# An *in vitro* assay of MCTS1-DENR-dependent re-initiation and ribosome profiling uncover the activity of MCTS2 and distinct function of eIF2D

**DOI:** 10.1101/2024.06.05.597545

**Authors:** Romane Meurs, Mara De Matos, Adrian Bothe, Nicolas Guex, Tobias Weber, Aurelio A. Teleman, Nenad Ban, David Gatfield

## Abstract

Ribosomes scanning from the mRNA 5′ cap to the start codon may initiate at upstream open reading frames (uORFs), decreasing protein biosynthesis. Termination at a uORF can lead to re-initiation, where the 40S subunit resumes scanning and initiates another translation event downstream. In mammals, the noncanonical translation factors MCTS1-DENR participate in re-initiation at specific uORFs, but knowledge of other *trans*-acting factors and uORF features influencing re-initiation is limited. Here, we describe a cell-free re-initiation assay using HeLa cell lysates. Comparing *in vivo* and *in vitro* re-initiation activities on uORF-containing model reporters, we validate that MCTS1-DENR-dependent re-initiation is accurately recapitulated *in vitro*. Using this system and ribosome profiling in cultured cells, we found that knockdown of the homolog eIF2D causes widespread gene expression deregulation unrelated to uORF translation, suggesting distinct functions from MCTS1-DENR. Additionally, we identified MCTS2, encoded by a retrogene copy of *Mcts1*, as an alternative DENR partner that promotes re-initiation *in vitro*, providing a plausible explanation for the striking clinical differences associated with *Denr* vs. *Mcts1* mutations in humans. Our findings on re-initiation and the new assay provide valuable insights and a powerful tool for future research on uORF features and *trans*-acting factors.

## Introduction

Upstream open reading frames (uORFs) are abundant, short regulatory elements found within the 5′ UTRs of an estimated 50 % of human transcripts [1]. uORF translation has long been recognised as a mechanism to tune the translation efficiency (TE) of the protein coding sequence (CDS) under physiological and stress conditions [2–5], and several cases of misregulated uORF-dependent translational control have been linked to human diseases including cancer [6–9]. Because uORFs intercept scanning ribosomes and reduce the probability that they engage in initiation downstream, they are generally inhibitory to protein biosynthesis from the CDS. Two main mechanisms are implicated in ensuring that CDS translation can take place nevertheless: first, in a process called leaky scanning, the 43S preinitiation complex can scan across the uORF start codon, thus ignoring the uORF; second, the ribosome can translate the uORF, terminate at its stop codon, and then engage in the process of re-initiation, where the small ribosomal subunit resumes scanning to a downstream start codon [1]. In contrast to canonical initiation, for which many implicated initiation factors and mechanistic details have been identified over the last decades, our understanding of re-initiation has remained incomplete. Thus, large knowledge gaps exist regarding the *trans*-acting protein machinery and to what extent it is shared with canonical initiation, and the *cis*-acting mRNA and uORF sequence requirements that predispose to re-initiation.

A peculiar case of re-initiation-competent uORFs that has been studied contains only a single amino acid – i.e., the start codon – followed immediately by a stop codon (termed ‘1-aa uORF’ or ‘start-stop uORF’ in the following) [10, 11]. In regular initiation, the initiator-tRNA is positioned in the ribosomal P-site, and on 1-aa uORFs the stop codon is thus already placed in the A-site, ready for termination and without the ribosome ever entering into elongation. Hence, and as shown for eIF3 [12], some canonical initiation factors from the uORF initiation event may likely still be available on the ribosome for subsequent re-initiation. *In vivo*, eIFs have been seen to remain associated with elongating ribosomes for roughly 12 codons, potentially enabling re-initiation after short uORFs but not long uORFs [13].

The heterodimeric, non-canonical initiation factor MCTS1-DENR has been identified as the first, re-initiation-specific translation factor with conserved activity in *Drosophila*, mouse and human [10, 11, 14–17]. *In vitro,* MCTS1-DENR shows the double activity of being able to recruit and release tRNA from the ribosomal P-site [14, 18]. The yeast orthologues, Tma20-Tma22, appear to mainly function in post-termination tRNA removal and 40S ribosome recycling rather than as promoters of re-initiation [19, 20]. In animal model systems, DENR loss-of-function followed by ribosome profiling has been used to identify CDS translation efficiency changes indicative of altered re-initiation, thus identifying likely direct MCTS1-DENR targets [10, 11, 17, 21]. These studies revealed that only a small fraction of all uORF-containing transcripts undergo MCTS1-DENR-dependent re-initiation and, while start-stop uORFs are highly represented among targets, longer uORFs were associated with MCTS1-DENR-dependent re-initiation as well. As a first hint towards the selectivity of MCTS1-DENR for certain uORFs, an enrichment for specific penultimate uORF codons was observed among targets in HeLa cells [21] and in yeast [20], leading to the hypothesis that MCTS1-DENR removes deacylated P-site tRNA from the terminated ribosome and that the efficiency of this process may be tRNA-specific.

The protein eIF2D is a close homologue of MCTS1-DENR, containing the same domains encoded on a single polypeptide. Like MCTS1-DENR, eIF2D (then still under the name LGTN, for Ligatin) was initially identified as a recycling and initiation factor based on its *in vitro* activities [14], and the yeast orthologue, Tma64, has been noted for its role in 40S subunit recycling, too [19]. Due to these similarities, eIF2D is frequently referred to as a re-initiation factor as well, and evidence for comparable functions of MCTS1-DENR and eIF2D on uORFs come from studies of *Atf4* regulation in *Drosophila* and in HeLa cells [21, 22]. However, eIF2D does not regulate an MCTS1-DENR-dependent synthetic model start-stop uORF *in vivo* [11], suggesting that the two factors may be active on distinct sets of uORFs and transcripts. Furthermore, it has recently been reported that the single knockout of yeast Tma64/*Eif2d* does not recapitulate the 40S recycling defect observed in the single knockout of Tma22/*Denr* [20]. Therefore, it would appear that the involvement of eIF2D in re-initiation and/or recycling still needs to be formally established.

In this study, we combined ribosome profiling, *in vivo* reporter assays and experiments using a cell-free translation system to investigate MCTS1-DENR-and eIF2D-dependent re-initiation. Using the *in vivo* and *in vitro* re-initiation assays, we measured the fluxes of ribosomes on the 5′ UTRs of two identified DENR-dependent model transcripts, *Klhdc8a* and *Asb8*, and addressed the effect of penultimate codon identity. Ribosome profiling from DENR and eIF2D loss-of-function cells pointed at different and distinct roles for the two factors in gene expression regulation and, importantly, indicated that eIF2D does not act as a general uORF re-initiation factor. Finally, we identified MCTS2 as an alternative DENR heterodimerisation partner *in vivo* whose activity in re-initiation we analysed and validated in our *in vitro* assay.

## Results

### Upstream open reading frame-containing transcripts *Klhdc8a* and *Asb8* are regulated by DENR *in vivo* and suitable for model reporter design

To gain insights into mechanism and targets of DENR-dependent re-initiation, we first identified uORF-containing transcripts for which CDS translation was altered after shRNA-mediated knock-down of *Denr* in murine NIH/3T3 cells (**Figure 1A**). To this end, we carried out ribosome profiling (Ribo-seq) and RNA-seq from *Denr* shRNA-treated cells, as well as from two different control-treated cells, exposed either to scramble (Scr) shRNA or to non-functional (nf) *Eif2d* shRNA that did not cause any knockdown (**Figure 1A**). From the obtained data, we calculated translation efficiencies (TEs, ratio CDS footprints to RNA) transcriptome-wide, thus identifying 223 and 6 transcripts, respectively, with significantly reduced and increased TE (applying FDR-corrected p < 0.1) (**Figure 1B**; Supplementary Table S1). The strong bias for reduced TEs was in line with our expectations for direct targets of a factor facilitating re-initiation on the CDS after uORF translation, and with previous findings [17, 21]; we further noted that the annotated 5′ UTRs of transcripts with reduced TE were significantly longer than would be expected from the transcriptome-wide distribution of 5′ UTR lengths (**Figure 1C**), compatible with increased uORF content. Indeed, annotation of translated uORFs from the footprint data showed strong enrichment among the DENR-responsive transcripts (**Figure 1D**). The lengths of the identified uORFs showed a broad distribution, with 1-aa/start-stop uORFs being the most abundant species but many longer, elongating uORFs found on the DENR-responsive transcripts as well (**Figure 1E**). We selected two transcripts to design re-initiation reporters, representing an elongating uORF (*Klhdc8a*, *Kelch domain containing 8A*; 13 amino acid uORF) and a start-stop uORF (*Asb8, Ankyrin repeat and SOCS box-containing 8*) (**Figure 1B**). Both mRNAs presented significantly reduced TE upon *Denr* knockdown (decrease by ∼70 % for *Klhdc8a* and by ∼30 % for *Asb8*; **Figure 1G**) and, moreover, their 5′ UTRs contained only one strongly translated, AUG-initiating uORF (**Figure 1F** and **Supplementary Figure S1A, B**) which is conserved in human and other mammals (**Supplementary Figure S1C, D**). We then confirmed that *Denr* KO in human HeLa cells indeed led to a strong downregulation of endogenous KLHDC8A protein as evaluated by immunoblot analysis using specific antibodies (**Figure 1J**). The lack of functional antibodies against mouse or human ASB8 precluded a similar analysis for this protein.

**Figure 1.**
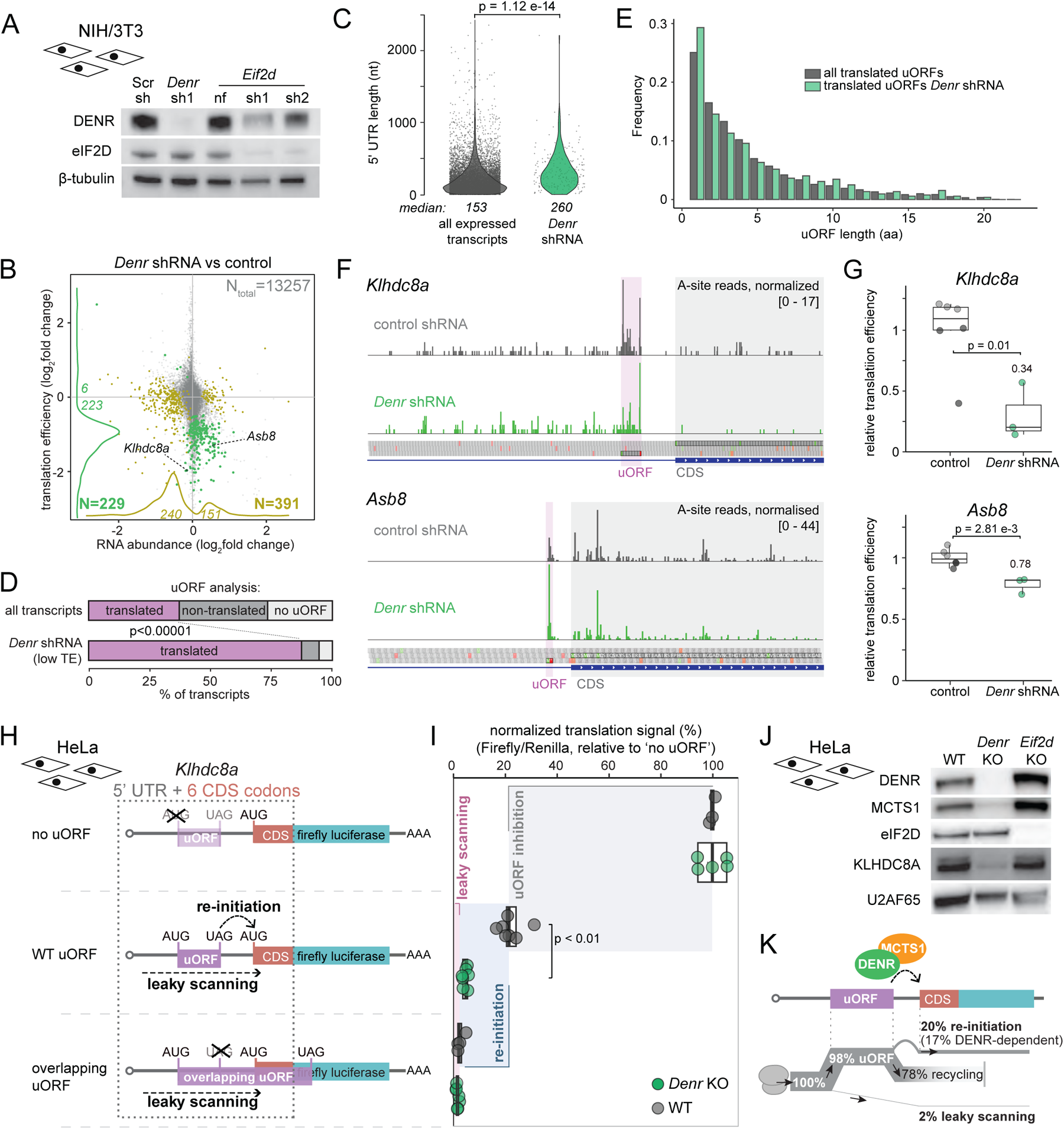
DENR is a re-initiation factor that promotes translation of hundreds of uORF-containing transcripts, including *Klhdc8a* and *Asb8*. **(A)** Immunoblot analysis of NIH/3T3 cells confirms DENR and eIF2D depletion upon lentiviral transduction with *Denr* or *Eif2d* shRNAs as compared to scramble (Scr) shRNA and *Eif2d* non-functional (nf) shRNA. β-tubulin serves as loading control. **(B)** Scatter plot of changes between *Denr* shRNA and control shRNA treated cells for RNA abundances vs. translation efficiencies (RPF counts/RNA counts), based on ribosome profiling and RNA-seq experiments. Evaluation is based on *Denr* shRNA (triplicates) and two independent control shRNAs (triplicates each). mRNAs with significant change for TE are represented in green and mRNAs with significant change for RNA abundance are represented in olive (adjusted p-values < 0.10, Wald test followed by FDR adjustment). Position of *Asb8* and *Klhdc8a* are indicated. **(C)** Violin plot showing 5′ UTR lengths of all expressed transcripts (n = 9203, grey) vs. transcripts with lower TE upon *Denr* depletion (n = 221, green). DENR-responsive genes have longer 5′ UTRs (median = 260 nt) than overall expressed transcripts (median = 153 nt) (p = 1.12e-14, Kolmogorov–Smirnov test). **(D)** Analysis of proportion of transcripts with at least one translated uORF. The transcripts with lower TE upon *Denr* depletion are enriched for translated uORFs. For all expressed transcripts (n = 9266), 3430 carry a translated uORF, whereas 3370 have a uORF sequence that is non-translated and 2465 have no uORF at all. In *Denr* shRNA cells, transcripts with significantly reduced TE (n = 179 after removal of transcripts with ambiguous/several expressed 5′ UTRs), 156 have translated uORFs vs. 13 non-translated uORFs and 10 no uORF (p-value < 1e-5, Fisher’s exact test). **(E)** The uORF length distribution of transcripts with reduced TE in *Denr* shRNA transduced cells (n = 522) is similar to that of all expressed transcripts (n = 7811) (p-value = 0.3717, Kolmogorov-Smirnov test). **(F)** Mapped footprint A-sites of *Klhdc8a* and *Asb8* transcripts in control and *Denr* knock-down cells. Read counts were normalized to library depth by subsampling and replicates were merged for increased coverage. **(G)** Quantification of translation efficiencies on the coding sequence for *Klhdc8a* and *Asb8* in cells treated with shRNAs targeting *Denr* (green) or controls (Scramble, Scr in dark grey; non-functional *Eif2d*-targeting shRNA, nf, in light grey); *Klhdc8a*: p = 0.01 and *Asb8*: p = 2.81e-3, two-tailed unpaired *t*-test. **(H)** Schematic of the *Klhdc8a* 5′ UTR reporters used for the *in vivo* and *in vitro* translation assays. The ‘no uORF reporter’ serves to evaluate the regulation by the 5′ UTR in the absence of a uORF (100 % signal). The ‘overlapping uORF reporter’ can only produce luciferase signal through leaky scanning. Comparison of the WT reporter, containing the relevant uORF, with the two former reporters allows to specifically calculate re-initiation. Not depicted: all plasmids also express Renilla luciferase for internal normalization. **(I)** Normalized luminescence signal (firefly/Renilla) of the *Klhdc8a* reporters after transduction in WT and *Denr* KO HeLa cells. The canonical initiation signal of the ‘no uORF reporter’ was set to 100 % in WT and *Denr* KO cells individually. The shaded boxes represent the signal that can be calculated for uORF inhibition (grey; ‘no uORF’ signal minus ‘WT uORF’ signal), re-initiation (blue; ‘WT uORF’ signal minus ‘overlapping uORF’ signal) and leaky scanning (pink; ‘overlapping uORF’ signal) (significance calculated using two-tailed unpaired *t*-test). **(J)** Western Blot analysis of WT, *Denr* KO and *Eif2d* KO HeLa cells validates depletion of KLHDC8A in the absence of DENR. U2AF65 serves as loading control. **(K)** Schematic representation of ribosomal fluxes on *Klhdc8a* 5′ UTR estimated from the results shown in panel (I).

We first used the *Klhdc8a* 5′ UTR (and the first codons of its CDS, fused in-frame to firefly luciferase) to establish a set of three different reporter genes that would allow us to estimate re-initiation and leaky scanning rates (**Figure 1H**). Thus, the *WT uORF* reporter contained the full, unmodified *Klhdc8a* 5′ UTR. Mutating the uORF start codon (*no uORF* reporter) allowed us to quantify luciferase signal when no inhibitory uORF is present. A third reporter contained a mutated uORF stop codon, which extended the uORF coding sequence all the way into the luciferase CDS, but in a different frame (*overlapping uORF* reporter). Thus, this reporter would give luciferase signal only through leaky scanning of ribosomes across the uORF start codon. After reporter delivery into HeLa cells (via lentiviral constructs that also contained a Renilla luciferase reporter for internal normalisation across samples), re-initiation signal can be extracted from the individual reporter readouts as shown in **Figure 1I**. The relative reduction of signal from *no uORF* to *WT uORF* reporters thus represents the overall inhibitory activity of the uORF, with remaining signal corresponding to the sum of leaky scanning-and re-initiation-mediated reporter translation. By further subtracting the *overlapping uORF* reporter signal that derives from leaky scanning, re-initiation is quantified. Using WT and *Denr* KO HeLa cells (**Figure 1J**), this assay indicated strong inhibition by the *Klhdc8a* uORF, and remaining signal that was largely DENR-dependent, as well as very low leaky scanning rates (**Figure 1I**). This data allowed us to propose a model of how ribosomal fluxes occur on the *Klhdc8a* reporter transcript *in vivo* (**Figure 1K**). Our analyses suggest that ∼98% of ribosomes initiate on the uORF and only ∼2% undergo leaky scanning. Of the uORF ribosomes about 19% re-initiate on the main CDS, a process that is largely DENR-dependent.

We concluded that translation of endogenous and reporter-encoded KLHDC8A was highly re-initiation-dependent and that our system based on three reporters, together with the use of WT vs. *Denr* KO cells, allowed the quantification of DENR-dependent re-initiation *in vivo*.

### An *in vitro* re-initiation assay recapitulates regulation and ribosome fluxes observed *in vivo*

Next, we set out to establish an *in vitro* assay that could recapitulate re-initiation. To this end, we prepared translation-competent extracts from HeLa cells, essentially adapting a recently reported protocol based on detergent-free cell lysis applying reproducible shearing forces via a dual centrifugation step [23] (**Figure 2A**). Under the gentle lysis conditions used, a sizeable proportion of cells remained intact (**Figure 2B**, pellet fraction), yet we found the supernatant to reproducibly recover highly translation-competent lysate that also contained the three proteins of interest for our study (MCTS1, DENR, eIF2D) (**Figure 2B**). Initial tests revealed that during the *in vitro* translation reaction, eIF2α became phosphorylated at Ser51 (**Figure 2C**), a modification that is associated with decreased initiation rates leading to low translational output. As previously reported [23, 24], the addition of recombinant GADD34Δ1-240 greatly counteracted Ser51 phosphorylation (**Figure 2C**). Indeed, in the absence of GADD34Δ1-240, luciferase signals that were obtained in *in vitro* translation assays for various *in vitro* transcribed, capped and polyadenylated reporter RNAs were >50 % reduced as compared to reporter signals in the presence of recombinant GADD34Δ1-240 (**Figure 2D**). As default condition in our experiments, we hence added recombinant GADD34Δ1-240 to all reactions. Finally, we further optimised the *Klhdc8a*-based reporters by shortening the 5′ UTR from its original ∼650 nt down to ∼180 nt, retaining the larger uORF environment yet removing far upstream 5′ UTR sequence (**Supplementary Figure S2A**). These short *Klhdc8a* 5′ UTR reporters showed strongly increased signals in our assays (**Supplementary Figure S2B**), in line with the notion that 5′ UTR length generally anticorrelates with CDS translation efficiency (e.g. [13, 25]).

**Figure 2.**
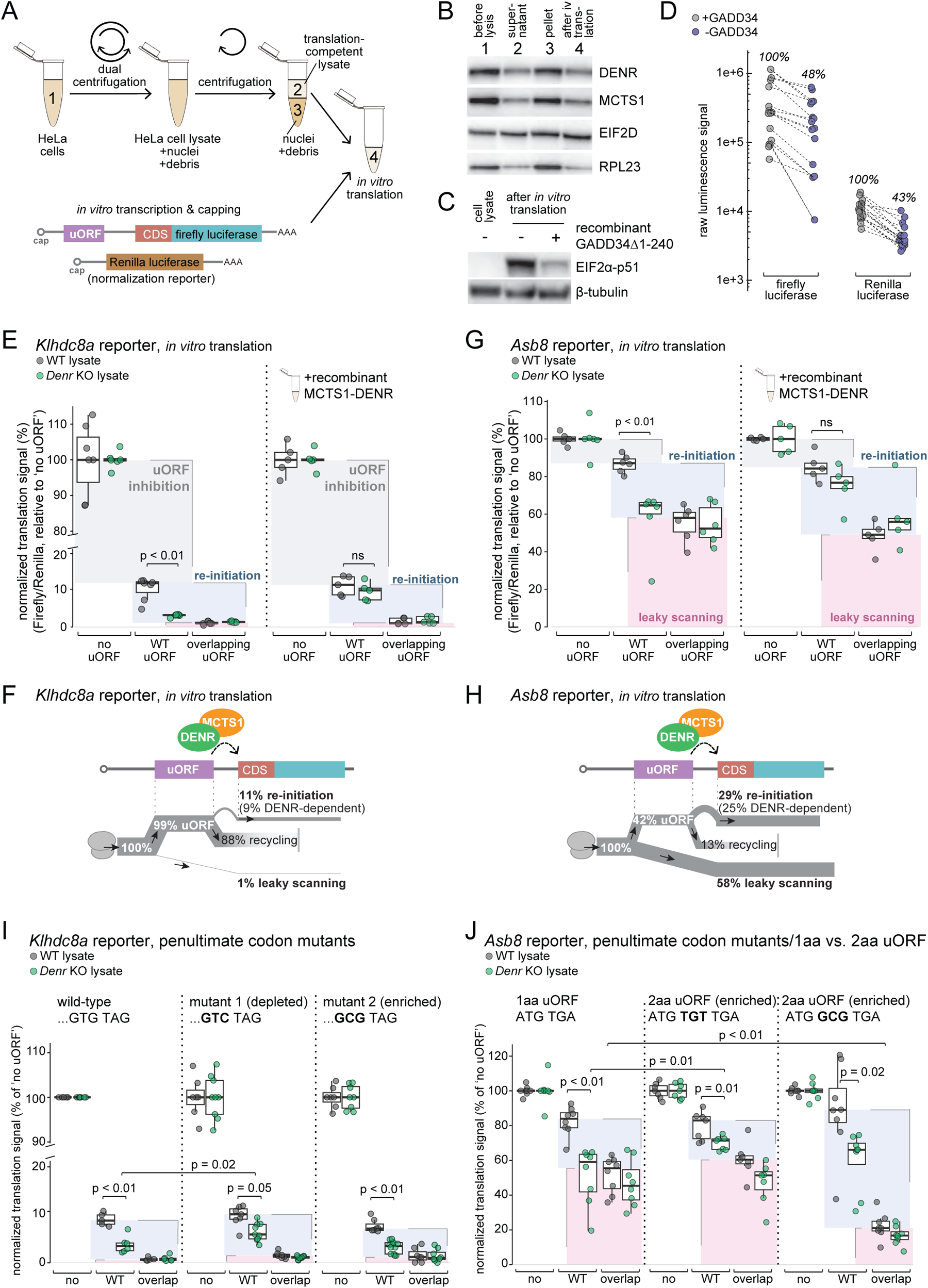
*In vitro* translation of *Klhdc8a* and *Asb8* reporters recapitulates regulation *in vivo*. **(A)** Outline of the preparation of HeLa translation-competent extracts and of the capped RNA reporters for the *in vitro* translation assays. The RNA for Renilla reporter was consistently used in all *in vitro* assays for normalization purposes. **(B)** Immunoblot analysis of proteins extracted from HeLa cells, i.e. whole cell lysate, supernatant and pellet post-dual-centrifugation and translation-competent extract post-translation. MCTS1, DENR, eIF2D and RPL23 are consistently recovered in the translation-competent extracts. **(C)** Immunoblot analysis of whole HeLa cell lysate and translation-competent extracts after *in vitro* translation reveals increased eIF2α-p51 during the *in vitro* translation reaction. β-tubulin serves as loading control. **(D)** Raw luminescence signals of firefly and Renilla luciferase reporters after *in vitro* translation in HeLa translation-competent extracts complemented with or without GADD34Δ1-240 validates reduced translation efficiency upon eIF2α phosphorylation. **(E)** Normalized luminescence signal (firefly/Renilla) of the *Klhdc8a* reporters after *in vitro* translation in WT and *Denr* KO HeLa lysates (left) and rescue assay with 0.5 µM recombinant MCTS1-DENR (right); ‘no uORF reporter’ signal was set to 100 % and uORF inhibition, re-initiation and leaky scanning were calculated as indicated by the shaded boxes and as in Fig. 1I (significance between ‘WT uORF’ signal in WT HeLa lysate and ‘WT uORF’ signal in *Denr* KO lysate calculated using two-tailed unpaired *t*-test). The lower and upper boundaries of the boxes correspond to the first and third quartiles while the middle line represents the median. The upper and lower whiskers extend to the largest and lowest values, limited to values that are not more distant than 1.5× the distance between the first and third quartiles. **(F)** Schematic representation of ribosomal fluxes on *Klhdc8a* 5′ UTR reporter estimated from the results shown in panel (E). **(G)** As in (E), but for the *Asb8* reporters. **(F)** As in (F) for the *Asb8* 5′ UTR in *vitro* data shown in (G). **(I)** Normalized luminescence signal (firefly/Renilla) of the *Klhdc8a* reporters with mutated uORF penultimate codon (GTG to GTC or GCG) reveals the effect of penultimate codon identity on DENR-dependence. **(J)** Normalized luminescence signal (firefly/Renilla) of the *Asb8* reporters with inserted codon (TGT or GCG) between the uORF start and stop codons indicates that lengthening of the uORF decreases DENR-dependence.

First, we carried out the *in vitro* translation assays on the set of *Klhdc8a* reporters (no uORF, WT uORF, overlapping uORF), using HeLa extracts prepared from both wild-type and *Denr* KO cells. These experiments revealed clear uORF-and DENR-dependence (**Figure 2E**, left side; identical outcomes for “long *Klhdc8a* 5′ UTR” reporter, see **Supplementary Figure S2C**), closely resembling our observations *in vivo* (**Figure 1I**). The calculated ribosomal fluxes showed excellent correspondence as well (**Figure 2F**). In particular, the uORF strongly impeded CDS expression with very low rate of leaky scanning (1%). Re-initiation-mediated reporter expression (11%) was quantitatively slightly lower than *in vivo* (19%, see **Figure 1K**), yet remained highly DENR-dependent (i.e., in *Denr* KO extracts the signal from the ‘WT uORF’ reporter was close to the leaky scanning signal from the ‘overlapping uORF’ reporter). Importantly, this lack of re-initiation in *Denr* KO extracts could be fully rescued through complementation with recombinant MCTS1-DENR (**Figure 2E**, right), confirming the direct involvement of these proteins in re-initiation *in vitro*. Note also that in *Denr* KO, MCTS1 is co-depleted (**Figure 1J**) as a result of its instability in the absence of its binding partner (as previously observed in [21, 26]), making a rescue with both proteins necessary. We concluded that our *in vitro* assays showed very good correspondence with re-initiation parameters observed *in vivo*.

Next, we assayed *in vitro* the start-stop uORF-containing *Asb8* reporter set (**Figure 2G**). These experiments revealed that this uORF was rather permissive to leaky scanning, likely due to poor Kozak context of its start codon; in particular, given that all stop codons (TGA, TAA, TAG) begin with a T nucleotide, start-stop uORFs by default have a (suboptimal) T in Kozak sequence +4 position. The lower efficiency of inhibition by the *Asb8* uORF (as compared to the *Klhdc8a* uORF) was in line with the translation efficiencies measured *in vivo* by Ribo-seq (**Figure 1G**). According to our calculations (**Figure 2H**), 58% of ribosomes underwent leaky scanning and 42% engaged in uORF translation. About two-thirds of uORF termination events led to re-initiation at the CDS and the remaining one-third did not (29% vs. 13%). Re-initiation was mostly MCTS1-DENR-dependent, as judged from the signals obtained in *Denr* KO extracts and from recombinant MCTS1-DENR rescue (**Figure 2G**).

MCTS1-DENR selectivity for certain uORFs has been linked to specific penultimate codons [19][21]. The *Klhdc8a* uORF terminates with the sequence ‘… GTG(Val) TAG(Stop)’, yet penultimate GTG was not observed as specifically associated with DENR-dependence previously [21]. In order to test penultimate codon influence in our assay, we tested reporters in which the GTG(Val) was mutated to GTC(Val) (among the reported significantly depleted penultimate codons in DENR-dependent uORFs [21]) and to GCG(Ala) (most strongly and significantly enriched in DENR-dependent uORFs according to [21]) (**Figure 2I**). We found that the inhibitory impact of the uORF and re-initiation rates were only weakly affected by these codon changes; in particular, re-initiation was lower (non-significant) than in wild-type codon constructs and it still remained strongly DENR-dependent. Interestingly, in the depleted penultimate codon case, GTC(Val), DENR-dependence of re-initiation was reduced, yet overall re-initiation capacity still intact. This observation is compatible with the idea that the depleted penultimate codon is able to partially decouple the re-initiation process from its dependence on these particular re-initiation factors.

Next, we examined *Asb8* reporter variants, converting the 1-aa uORF to a 2-aa uORF (and thus from start-stop to elongating uORF) and testing different penultimate codons. Thus, we inserted either a TGT(Cys), which is (according to [21]) the only significantly enriched penultimate codon starting with ‘T’ (hence preserving the +4 Kozak context of the start codon), or GCG(Ala), which is a highly enriched codon and additionally improves the Kozak context for uORF initiation (**Figure 2J**). In the *in vitro* translation assays, GCG insertion indeed led to strongly decreased leaky scanning rates, converting the weakly translated 1-aa uORF to a strongly translated 2-aa uORF (**Figure 2J**, right), compatible with the improved Kozak consensus associated with the +4 position ‘G’. Lengthening the uORF with TGT(Cys) had little consequences on leaky scanning and overall re-initiation rate, yet DENR-dependence appeared reduced (**Figure 2J**, middle). We concluded that re-initiation rates were not *per se* different between start-stop and elongating uORFs, but the DENR-dependence of re-initiation was modulated, which is in agreement with the reported preferential MCTS1-DENR regulation of start-stop uORFs [11, 21].

### Phospho-eIF2α levels only mildly affect re-initiation rates of *Klhdc8a* and *Asb8* reporters

Phosphorylation of eIF2α is key to the integrated stress response (ISR) that lowers bulk translation initiation and promotes preferential translation of stress-related mRNAs through a mechanism that involves uORFs, re-initiation and start codon selection [27]. The current model deduced from studying transcripts such as *Atf4/Gcn4* suggests that after uORF termination, the re-scanning 40S subunit becomes rapidly re-initiation-competent when non-phosphorylated (active) eIF2α is abundant. This leads to translation of a relatively close (87 nt) downstream, second uORF rather than the CDS that is more distant (184 nt). However, when p-eIF2α levels are high, the probability of recruiting active eIF2α in time for the second uORF is lower, and re-initiation will more often occur further downstream, on the CDS. Recent work has highlighted the importance of both MCTS1-DENR and eIF2D for *Atf4* CDS translation [21, 22]. Despite these links, whether eIF2α activity is a rate-limiting factor for MCTS1-DENR-mediated re-initiation *per se*, and to what extent the reported effects are a consequence of global initiation decrease, remains unclear. We reasoned that our well-controlled *in vitro* system would allow us to address such questions.

We thus carried out our assays in the presence vs. absence of GADD34Δ1-240, which led to low and high Ser51 phosphorylation levels of eIF2α, respectively (**Figure 2C**). The level of Ser51 phosphorylation occurring in the absence of GADD34Δ1-240 was even higher than that seen after a standard ISR induction protocol by tunicamycin treatment of cells (**Supplementary Figure S3A**). For the *Klhdc8a* reporters, absolute signals were significantly reduced in the absence of GADD34Δ1-240 as expected (analogous to experiments in **Figure 2D**), yet after normalisation to the (likewise reduced) Renilla luciferase signal, the observed levels of leaky scanning (slightly up) and uORF-mediated inhibition (slightly down), the re-initiation rate (slightly down) and its DENR-dependence (up), and the calculated ribosomal fluxes were highly similar between both conditions (**Figure 3A, B**). We concluded that in the *Klhdc8a* reporter *in vitro* experiments, the availability of non-phosphorylated eIF2α was not *per se* a strongly rate-limiting step for re-initiation. On the *Asb8* reporter – where the intercistronic distance between uORF and CDS is very short (17 nt, as compared to 73 nt in *Klhdc8a*) – we found overall more variability in the experimental outcomes in the absence of GADD34Δ1-240 (**Figure 3C, D**). We observed a tendency to increased leaky scanning and reduced re-initiation (both non-significant). In an *in vivo* experiment on the *Klhdc8a* reporter set, combined with tunicamycin treatment, we also observed a weak reduction in re-initiation rate (**Supplementary Figure S3B, C**) and validated that under these conditions the translation of positive control reporters carrying the 5′ UTRs of *Atf4* and *Atf5* upstream of luciferase were indeed upregulated (**Supplementary Figure S3D**). Taken together, these results suggest that eIF2α levels are only moderately limiting for re-initiation.

**Figure 3.**
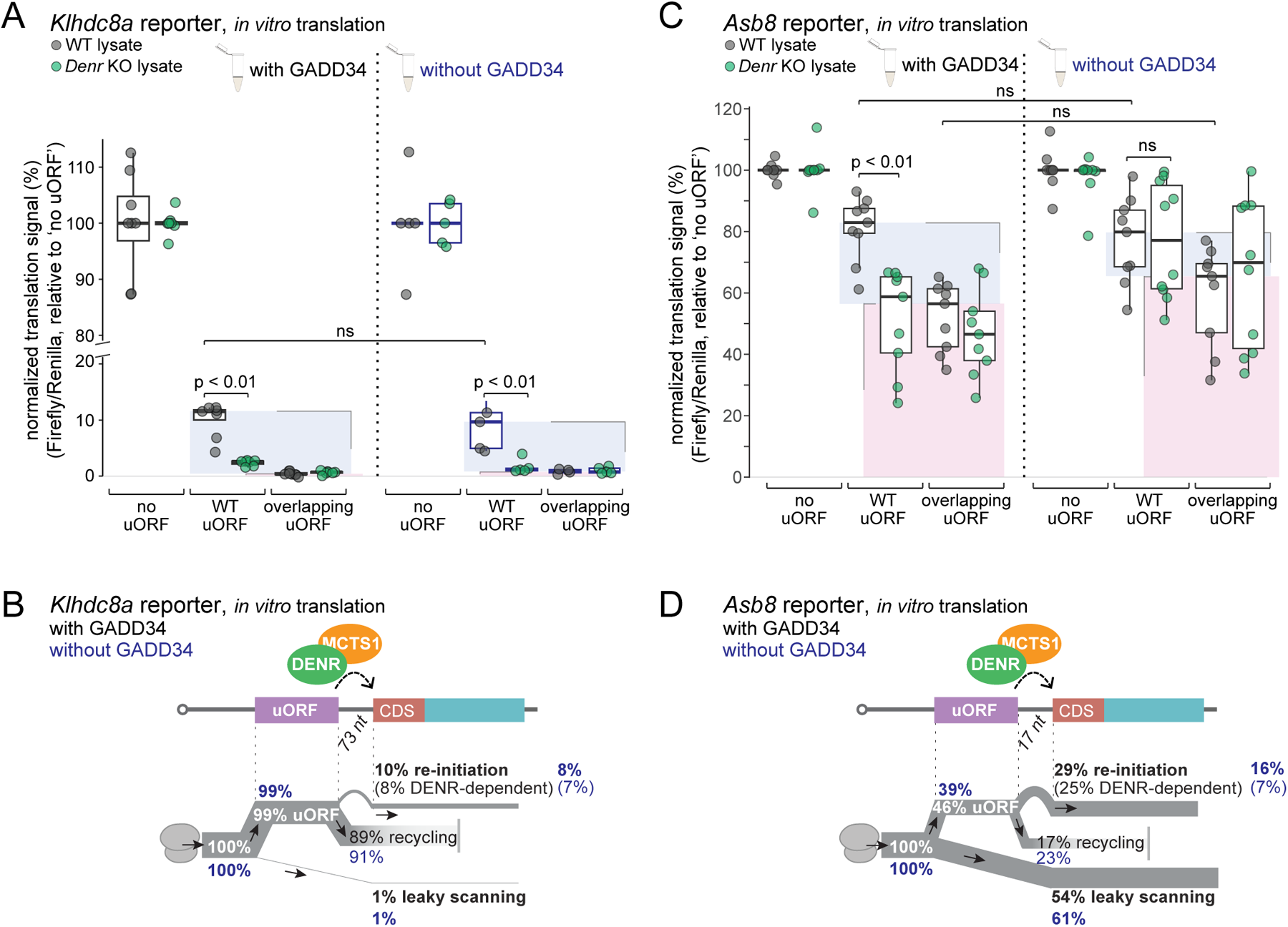
eIF2α phosphorylation status moderately affects re-initiation rate of start-stop uORF and longer uORF reporters. **(A)** Normalized luminescence signal of *Klhdc8a* reporters after *in vitro* translation in WT and *Denr* KO HeLa extracts complemented with 16 ng/μl recombinant GADD34Δ1-240 (left) or without GADD34Δ1-240 (right) (significance calculated using two-tailed unpaired *t*-test). **(B)** Schematic representation of ribosomal fluxes on *Klhdc8a* 5′ UTR *in vitro* estimated from the results shown in panel (A). Quantifications in presence of GADD34Δ1-240 are shown in black and without GADD34Δ1-240 in blue. **(C)** As in (A) for *Asb8* reporters. **(D)** As in (B) for *Asb8* reporter data shown in (C).

### Limited evidence for a role of eIF2D in re-initiation *in vivo* and *in vitro*, pointing to uORF-independent cellular functions in gene expression regulation

Next, we extended our analyses to the close MCTS1-DENR homologue, eIF2D, which contains the same domains encoded as a single polypeptide, linked by an additional WH domain (**Figure 4A**). The high similarity between the proteins, in addition to the findings that the yeast orthologues show shared activities, has led to the assumption that the homology in mammals extends to their function. To test this hypothesis, we downregulated eIF2D using specific shRNAs (**Figure 1A**) and carried out Ribo-seq and RNA-seq identically to the experiment in *Denr*-depleted cells shown in **Figure 1B**. For a first comparison between the two genotypes, i.e. *Eif2d*-depleted and *Denr*-depleted, we evaluated in a transcriptome-wide fashion the relative ribosome distribution between mRNA 5′ UTR and CDS sequences. This measure is expected to increase when a re-initiation factor is depleted because ribosomes are selectively lost from CDS sequences but will still translate uORFs when re-initiation in inhibited [17]. Indeed, the distribution of the changes seen for 5′ UTR-to-CDS footprint ratios between *Denr* shRNA-treated cells vs. control cells was strongly shifted to the positive (**Figure 4B**), indicating globally detectable ribosome redistribution in the absence of MCTS1-DENR, consistent with previous findings [17]. By contrast, the shift was considerably smaller in the cells that had been treated with *Eif2d* shRNAs (**Figure 4B**). Next, we carried out an analysis identical to that applied to *Denr* knockdown cells in **Figure 1B**, to determine the effects induced by *Eif2d* depletion at the level of CDS translation efficiencies (**Figure 4C**). This analysis, too, revealed strikingly different transcriptome-wide effects between the homologues. We thus detected much fewer transcripts with a decreased TE upon *Eif2d* knockdown (n = 78, with FDR-corrected p < 0.1; Supplementary Table S2) (**Figure 4C**) as compared to *Denr* knockdown cells (n = 223) (**Figure 1C**), and the overlap between the two sets was minimal (**Supplementary Figure S4A, B**). Moreover, the mRNAs with low TE in *Eif2d*-depleted cells tended to have short 5′ UTRs (**Supplementary Figure S4C**) and they were not enriched for translated uORFs (**Figure 4D**). We also noted that many of the translationally affected transcripts were highly abundant, including >25 mRNAs encoding for ribosomal proteins and general translation machinery (**Supplementary Figure S4D, E**). Particularly strong consequences of eIF2D loss-of-function were evident for RNA abundances, with a large number of mRNAs significantly up (n = 460) and downregulated (n = 617) (**Figure 4C**) and, here, abundant transcripts were overrepresented as well (**Supplementary Figure S4F**) and associated with cellular processes such as the cell cycle and translation (**Supplementary Figure S4G, H**). Alterations to the cell cycle could indeed be further validated as a robust phenotype following eIF2D loss-of-function (**Supplementary Figure S4I**). Taken together, these findings suggested that, first, the knockdown of *Eif2d* was phenotypically highly consequential as judged by the widespread gene expression reprogramming that it induced. However, second, it would seem that the primary activity of eIF2D may not lie in regulating the re-initiation of uORF-containing transcripts.

**Figure 4.**
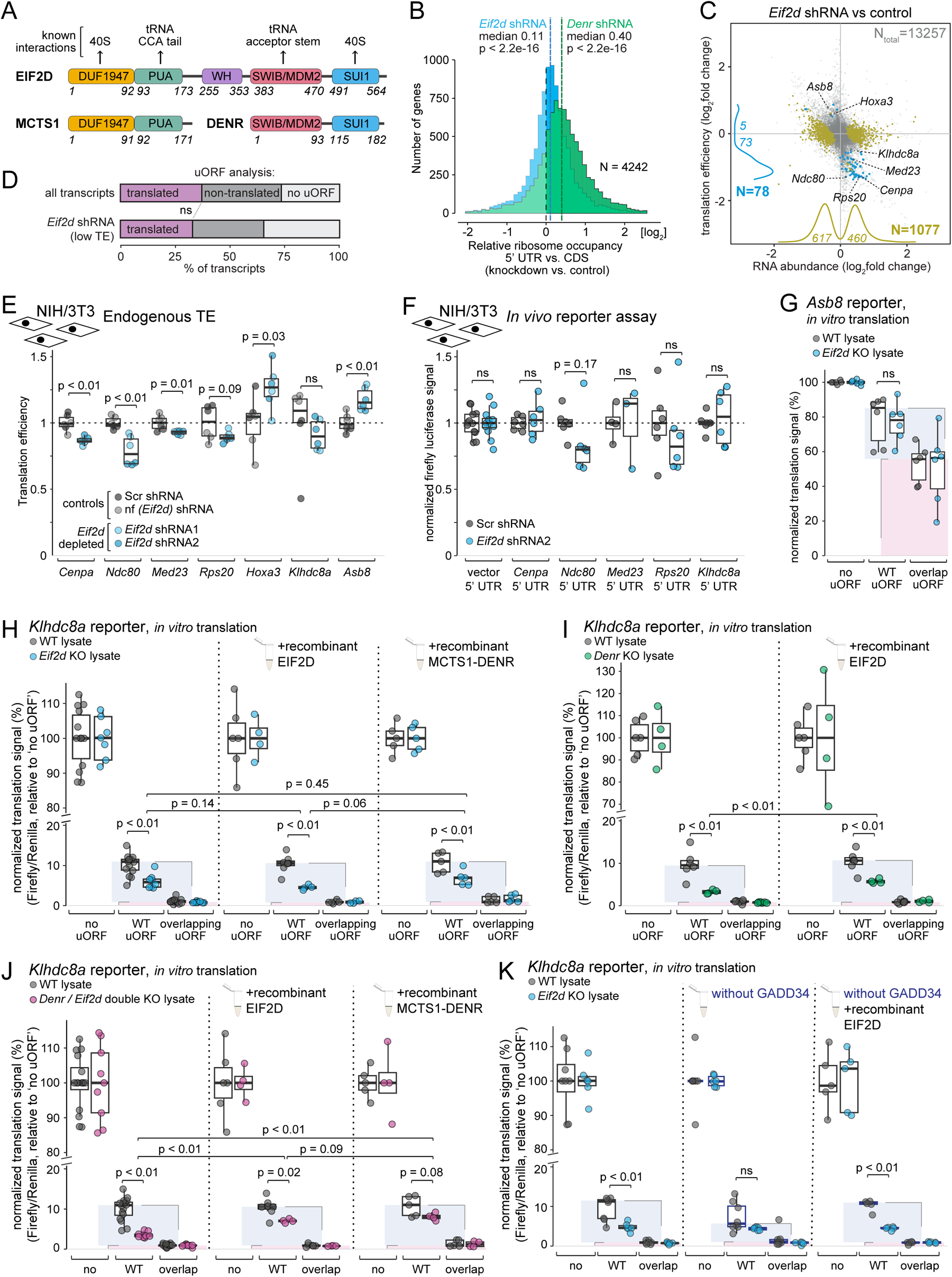
No evidence for eIF2D uORF re-initiation activity *in vivo* but low activity *in vitro*. **(A)** Schematic representation of human MCTS1-DENR and eIF2D proteins, with domains indicated. The relevant amino acids positions defining the domains are specified. **(B)** Distribution of ribosomes on 5′ UTR relative to CDS in *Denr* knock-down (in green) or *Eif2d* knock-down (in blue) vs. control cells (n = 4242). *Denr* knock-down cells show a strong redistribution of ribosomes from CDS to 5′ UTR whereas the effect is milder but still significant for *Eif2d* knock-down cells (p-values calculated using Wilcoxon signed rank test). **(C)** Scatter plot of changes between *Eif2d* shRNA and control cells for RNA abundances vs. translation efficiencies (RPF counts/RNA counts), based on ribosome profiling and RNA-seq experiments. Evaluation is based on two independent *Eif2d* shRNAs (triplicates each) and two independent control shRNAs (triplicates each). mRNAs with significant change on TE were represented in blue and mRNAs with significant change on RNA abundance were represented olive green (adjusted p-values < 0.10, Wald test followed by FDR adjustment). Positions of *Asb8*, *Klhdc8a*, *Hoxa3*, *Med23*, *Rps20*, *Ndc80* and *Cenpa* are indicated. **(D)** Analysis of proportion of transcripts with at least one translated uORF. The transcripts with lower TE upon *Eif2d* depletion are not enriched for translated uORFs. For all expressed transcripts (n = 9266), 3430 carry a translated uORF, whereas 3370 have a uORF sequence that is non-translated and 2465 have no uORF at all. In *Eif2d* shRNA cells, transcripts with significantly reduced TE (n = 58 after removal of transcripts with ambiguous/several expressed 5′ UTRs), 19 have translated uORFs vs. 19 non-translated uORFs and 20 no uORF (p-value = 0.59, Fisher’s exact test). **(E)** Quantification of translation efficiencies on the coding sequence of selected transcripts with lower TE upon *Eif2d* depletion in control and *Eif2d*-depleted cells (p-values calculated using a two-tailed unpaired *t*-test). **(F)** Normalized luminescence signal (firefly/Renilla) of lentivirally transduced reporters with 5′ UTRs of selected transcripts with lower TE upon *Eif2d* depletion (significance calculated using two-tailed unpaired *t*-test). The signal of individual reporter expressed in WT cells was set to 100 %. **(G)** Normalized luminescence signal (firefly/Renilla) of the *Asb8* reporters after *in vitro* translation in WT and *Eif2d* KO HeLa extracts (p-value calculated using two-tailed unpaired *t*-test). **(H)** Normalized luminescence signal (firefly/Renilla) of the *Klhdc8a* reporters after *in vitro* translation in WT and *Eif2d* KO HeLa extracts (left), supplemented with 0.5 µM recombinant eIF2D (middle) or supplemented with 0.5 µM of recombinant MCTS1-DENR (right, p-values calculated using two-tailed unpaired *t*-test). **(I)** Normalized luminescence signal (firefly/Renilla) of the *Klhdc8a* reporters after *in vitro* translation in WT and *Denr* KO HeLa extracts (left) or supplemented with 0.5 µM recombinant eIF2D (right); p-value calculated using two-tailed unpaired *t*-test. **(J)** Normalized luminescence signal (firefly/Renilla) of the *Klhdc8a* reporters after *in vitro* translation in WT and *Denr / Eif2d* double KO HeLa extracts (left), supplemented with 0.5 µM recombinant eIF2D (middle) or supplemented with 0.5 µM recombinant MCTS1-DENR (right); p-values calculated using two-tailed unpaired *t*-test. **(K)** Normalized luminescence signal (firefly/Renilla) of the *Klhdc8a* reporters after *in vitro* translation in WT and *Eif2d* KO HeLa extracts treated with GADD34Δ1-240 (left), without GADD34Δ1-240 (middle), or without GADD34Δ1-240 and complemented with 0.5 µM eIF2D (right); p-values calculated using two-tailed unpaired *t-*test.

To further investigate the above observations on potential eIF2D targets, we selected several candidate transcripts to test if their TE regulation was 5′ UTR-mediated. Of these, *Cenpa*, *Ndc80*, *Med23* and *Rps20* were picked as candidate mRNAs with low TE from **Figure 4C**. We also included *Hoxa3*, which has been identified as a *bona fide* eIF2D-regulated mRNA in a previous study [28]. According to the published findings, eIF2D is implicated in uORF-mediated inhibition, rather than in re-initiation, of *Hoxa3* CDS translation. Indeed, our *in vivo* data confirmed *Hoxa3* CDS TE up-regulation upon *Eif2d* depletion (**Figure 4E**; note that *Hoxa3* is very lowly expressed in our cells and did not feature among the significantly changed TE transcripts in **Figure 4C**, but the differences were statistically significant in the direct comparison of TEs shown in **Figure 4E**). *In vivo* luciferase reporter assays using the 5′ UTRs of the selected endogenous candidate transcripts did not reveal significant regulation by eIF2D (**Figure 4F**). We also analysed *Klhdc8*a (endogenous and 5′ UTR reporter), but we found no indication that eIF2D was involved in its re-initiation *in vivo* (**Figure 4E, F**). Endogenous *Asb8* showed an increase in translation efficiency upon *Eif2d* depletion (**Figure 4E**), contrasting the effect seen upon *Denr* depletion.

We next tested the outcome of eIF2D loss-of-function and rescue in our *in vitro* assay. In lysates from *Eif2d* KO HeLa cells (**Figure 1J**), no difference in re-initiation activity or leaky scanning was detectable for the *Asb8* reporter (**Figure 4G**). By contrast, for the *Klhdc8a* reporter, re-initiation rates in *Eif2d* KO lysates were reduced by about half (**Figure 4H**). However, neither recombinant eIF2D nor recombinant MCTS1-DENR were able to fully rescue the low signal obtained with the WT uORF reporter in *Eif2d* KO lysates (**Figure 4H**). These observations were most compatible with the model that lower re-initiation signal was an indirect effect of eIF2D loss, e.g. due to secondary effects related to the widespread gene expression reprogramming seen in **Figure 4C**. Thus, we would not expect indirect effects to be rescuable *in vitro*, whereas a direct role in the re-initiation process itself should be rescued by adding the needed re-initiation factor as a recombinant protein. In *Eif2d* knockout extracts, MCTS1-DENR is abundant and may mask the re-initiation function of eIF2D. To address this possibility, we carried out two experiments. First, using re-initiation-deficient *Denr* KO extracts and recombinant eIF2D, we were indeed able to partially rescue re-initiation activity (**Figure 4I**), although not as efficiently as with recombinant MCTS1-DENR (compare with **Figure 2E**). Second, we used *Denr + Eif2d* double knockout cell lysates. *In vitro* translation reactions in these extracts showed reduced re-initiation activity (**Figure 4J**, left) that was very similar to *Denr* single knockouts (**Figure 2E**). Addition of recombinant MCTS1-DENR showed the expected rescue (**Figure 4J**, right) and, interestingly, also recombinant eIF2D partially rescued under in these conditions, i.e. in the absence of MCTS1-DENR (**Figure 4J**, middle). Taken together, these results indicate that eIF2D is likely not an essential component of the re-initiation machinery *in vivo* (**Figure 4C**) and *in vitro* (**Figure 4H**). However, when MCTS1-DENR is absent, adding high concentrations of eIF2D (0.5 μM) can partially compensate for MCTS1-DENR loss. Thus, it is a probable scenario that eIF2D has in principle retained some shared activity with MCTS1-DENR, but its main cellular functions seem to lie outside of uORF re-initiation.

Finally, it is also possible that eIF2D is active in re-initiation in a conditional fashion; in particular upon induction of the integrated stress response that leads to unavailability of eIF2α, eIF2D may be able to deliver initiator-tRNA to the 40S ribosome. We addressed this possibility by carrying out *in vitro* translation in *Eif2d* knockout extracts (without/with recombinant eIF2D rescue) in the absence of GADD34Δ1-240 (**Figure 4K**). Also under these conditions, we observed the (most likely) indirect/secondary effects of eIF2D loss that could not be rescued by recombinant eIF2D, and the experiment revealed no further indications for a specific re-initiation role of eIF2D.

### Identification of MCTS2 as a functional DENR heterodimerization partner

It has been observed that MCTS1 and DENR are mutually dependent on each other for protein stability [21, 26], giving rise to the idea that they act as obligate heterodimers. However, closer inspection of the available cell culture data would suggest that the depletion of MCTS1 in *Denr* KO cells is stronger than the depletion of DENR in *Mcts1* KOs [21]; moreover, *in vivo* phenotypes in mouse neuronal migration are stronger upon DENR loss than MCTS1 loss [29]. These findings could indicate that DENR is indeed an obligatory component of the heterodimer, but that it can interact with and be stabilised by alternative heterodimerization partners as well. To obtain insights into this hypothesis, we sought to purify DENR-interacting proteins by a co-immunoprecipitation and mass-spectrometry approach. Using CRISPR gene editing, we modified the endogenous *Denr* locus in NIH/3T3 cells such that it expressed DENR protein with a C-terminal tag suitable for immunoprecipitation via a 3×FLAG sequence as well as for degron-mediated protein depletion via the dTAG / dTAG-13 degrader system [30] (**Figure 5A**). Genotyping indicated successful modification of both alleles in selected cell clones (**Figure 5B**) and immunoblot analysis validated the functionality of the degron system (**Figure 5C**). We then analysed by mass-spectrometry the proteins enriched in FLAG-IP performed on extracts from control-treated vs. dTAG-13-treated cells (**Figure 5D**). The two most strongly enriched co-purifying proteins were MCTS1 and MCTS2. This finding indicates that DENR assembles into two distinct heterodimeric complexes, MCTS1-DENR and MCTS2-DENR. *Mcts2* is a paralog and retrogene copy of *Mcts1* that has been noted for its transcription from an imprinted gene region in mouse and humans, with predominant expression from the paternal allele [31]. A recent study on human *Mcts1* deficiency and associated disease phenotypes has speculated on redundancy with *Mcts2*, yet based on reporter assays and other analyses, it came to the conclusion that this paralog was largely non-functional [32]. By contrast, our interaction data collected from endogenous proteins suggests that both MCTS paralogs form *bona fide* heterodimers with DENR. The *Mcts1* and *Mcts2* coding sequences differ at 81 individual nucleotide positions that are distributed quite evenly across the overall CDS of 546 nt (**Supplementary Figure S5A**), allowing to unambiguously assign RNA-seq reads and Ribo-seq footprints to the two paralogous sequences. We could thus compare the expression of the *Mcts* paralogs, which indicated higher levels of *Mcts2* than *Mcts1* in our cells (**Figure 5E**). These findings strongly indicate that MCTS2 is a relevant interaction partner of DENR besides MCTS1.

**Figure 5.**
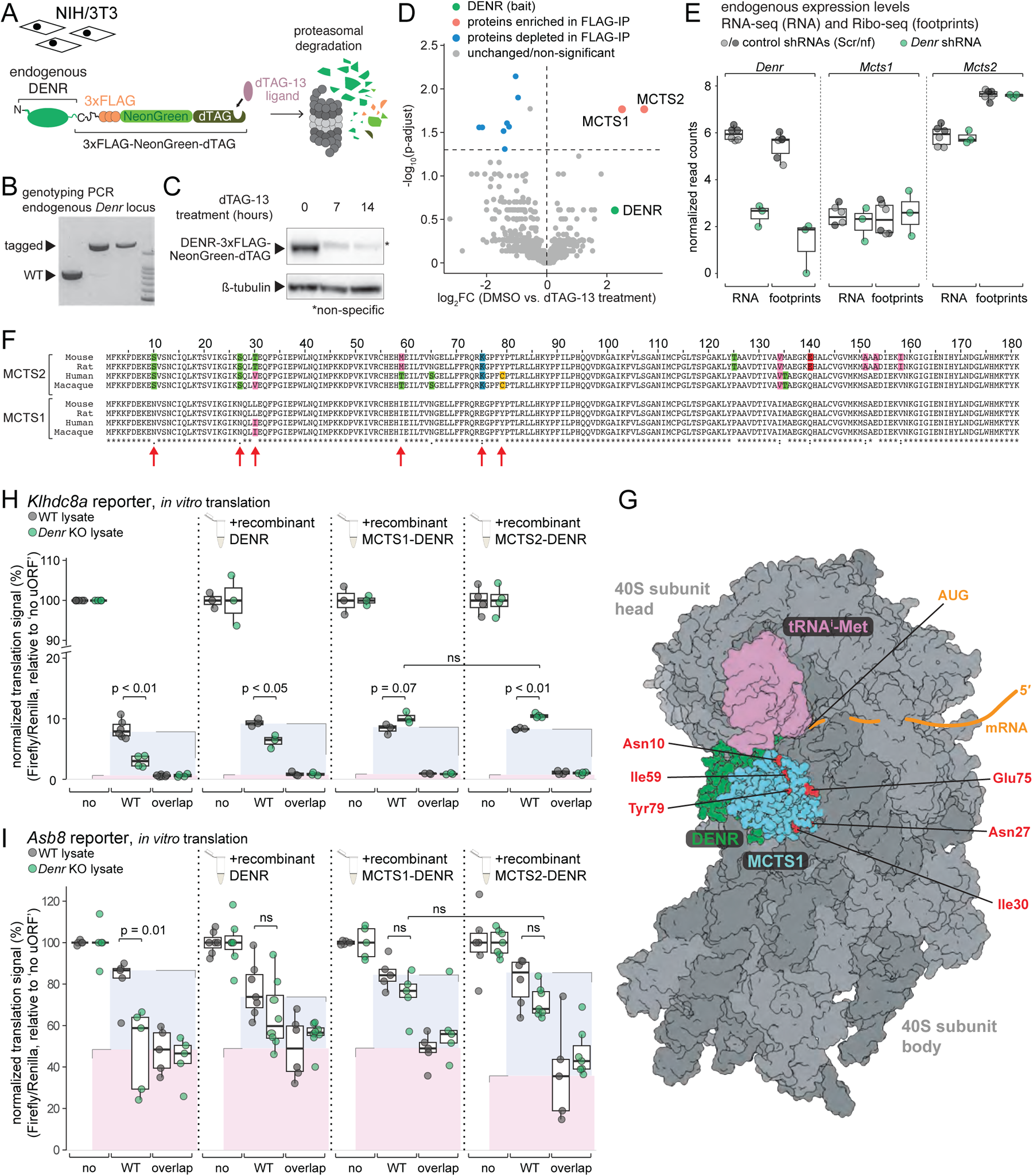
MCTS2 interacts with DENR *in vivo* and promotes re-initiation *in vitro*. **(A)** Schematic representation of endogenous DENR tagged with 3×FLAG-NeonGreen-dTAG. Treatment with the dTAG-13 ligand targets DENR-3×FLAG-NeonGreen-dTAG to proteasomal degradation. **(B)** Genotyping PCR analysis of *Denr*-tagged clones validates the insertion of the 3×FLAG-NeonGreen-dTAG cassette upstream of *Denr* stop codon. Tagged *Denr* amplicon has a length of 2067 bp and WT *Denr* of 885 bp. **(C)** Western Blot analysis of NIH/3T3 DENR-3×FLAG-NeonGreen-dTAG cells treated for 7 or 14 h with 500 nM dTAG-13 shows complete depletion of DENR upon treatment. Asterisk indicates a non-specific band that migrates at a similar height of the tagged protein. **(D)** Mass spectrometry analysis of proteins co-immunoprecipitated with endogenously tagged DENR in non-treated vs. dTAG-13-treated cells reveals the interaction of MCTS2 with DENR. Only DENR and significant hits enriched in non-treated cells are shown (p-values calculated using two-tailed unpaired *t*-test followed by correction for multiple testing). **(E)** Normalized RNA abundances and ribosome footprints of *Denr*, *Mcts1* and *Mcts2* in control and *Denr* depleted NIH/3T3 cells (p-values calculated using two-tailed unpaired *t*-test). **(F)** Alignment of human, rhesus macaque, mouse and rat MCTS1 and MCTS2 amino acid sequences. The amino acids changing relative to mouse MCTS1 are highlighted with colors according to their side-chain chemistry. **(G)** Structural model of MCTS1-DENR (turquoise/green) in interaction with the 40S ribosomal subunit (grey), Met-tRNA^Met^_i_ (magenta) and an mRNA shown schematically (orange). The amino acids that undergo changes in MCTS2 are indicated in red in the structure and labeled. PDB accession numbers used for depiction: 5vyc (40S and MCTS1-DENR), 5oa3 (initiator tRNA), 6ms4 (MCTS1-interacting domain of DENR) **(H)** Normalized luminescence signal (firefly/Renilla) of the *Klhdc8a* reporters after *in vitro* translation in WT and *Denr* KO HeLa lysates (left), substituted with 0.5 µM of recombinant DENR, MCTS1-DENR or MCTS2-DENR as indicated (p-values calculated using two-tailed unpaired *t-*test). **(I)** As in (H), for the *Asb8* reporters.

Human and mouse MCTS1 protein sequences are identical apart from one leucine to isoleucine substitution at amino acid position 30 (**Figure 5F**). In MCTS2, divergence to MCTS1 can be found at 14 positions when aligning mouse, rat, human and macaque protein sequences, with 4 substitutions common to MCTS2 across species and other changes specific to the analysed rodents or primates (**Figure 5F**). Using available structural data from *in vitro* reconstituted MCTS1-DENR in complex with the 40S ribosome, initiator-tRNA and a viral model RNA [33], we found that MCTS2-specific amino acid changes affected positions that cluster all in the same region of the fold but outside of the interaction surface between MCTS1-DENR and also outside of that between MCTS1-DENR and the 40S ribosomal subunit (**Figure 5G**). Hence, the capacity of MCTS2 vs. MCTS1 to interact with DENR, and of MCTS2-DENR vs. MCTS1-DENR to interact with 40S can be expected to be comparable. However, the substitutions – of which in particular the change at amino acid position 75 from glutamate (negative charge) in MCTS1 to lysine (positive charge) in MCTS2 (**Figure 5F**) would have the potential to significantly alter protein properties – could affect additional, so far unknown interactions taking place at this solvent-exposed surface within the complex (**Figure 5G**). To gain insights into whether MCTS1-DENR and MCTS2-DENR could redundantly act in re-initiation, we tested both as recombinant proteins on *Klhdc8a* and *Asb8* reporters. These experiments revealed that both paralogs were able to rescue *Denr* KO extracts in a comparable fashion (**Figure 5H, I**). Taken together, we concluded that *Mcts2*, although previously categorised as a pseudogene [34], is expressed to relevant levels and produces protein that assembles with DENR into protein complexes *in vivo*. *In vitro*, the paralogous complexes show similar activity on our two re-initiation model substrates, indicating redundancy in activity at least for certain substrates.

## Discussion

Translation re-initiation is a process that is at odds with several principles of canonical initiation, termination and post-termination subunit recycling and that remains mechanistically particularly poorly understood. The MCTS1-DENR heterodimer is so far the only reported *trans*-acting factor that is specific to re-initiation, i.e. it does not appear to be essential for general translation as well – quite in contrast to several other proteins that have also been linked to re-initiation in mammals, such as components of eIF3 [13, 35, 36] or eIF4 [13]. For these general eIFs, the analysis of specific re-initiation effects *in vivo*, for example through loss-of-function experiments, is challenging, as they cannot easily be deconvoluted from the abundant global translational alterations and their secondary effects. A defined *in vitro* assay that faithfully recapitulates and quantifies re-initiation gives the possibility of dissecting this process specifically, allowing for targeted, better interpretable experiments. Therefore, we view the development of the *in vitro* re-initiation assay presented in this study as an important methodological advance that will allow the characterisation of *cis*-acting mRNA/uORF sequence requirements and *trans*-acting protein machinery under defined conditions. Our careful benchmarking of the assay, including the use of different reporter constructs and variants, the demonstration of its dependence on MCTS1-DENR using knockout lysates and rescue with recombinant proteins, and the calculation of ribosomal fluxes, demonstrate the suitability of our system for detailed future mechanistic dissection of re-initiation. For example, a strategy to test the requirement for general eIFs in re-initiation could consist in producing HeLa *in vitro* translation lysates devoid of the protein of interest (e.g. through acute degron-mediated depletion, just before harvest of the HeLa cells), and to drive uORF translation on the *Klhdc8a* reporter through IRES sequences (e.g. the “factorless” Cricket Paralysis Virus IRES [37]) that bypass the requirement for the particular eIF of interest. Thus, it should be possible to query eIF requirement specifically in re-initiation. Other attractive future applications for our *in vitro* assay include structural biology approaches (i.e., cryo-EM on re-initiation complexes) or compound screens (i.e., testing for specific inhibitors of re-initiation).

In addition to the methodological advancement, our study provides important new biological insights, notably regarding MCTS1-DENR and the paralogous MCTS2, as well as on the role that eIF2D plays in gene expression regulation. In particular our observations on eIF2D are surprising, as the *in vivo* and *in vitro* evidence that we provide indicates that the main functions of this protein may not lie in uORF re-initiation after all. In retrospect, the case of eIF2D carrying re-initiation activity in mammals *in vivo* was always heavily built on circumstantial evidence. This evidence was, mainly, the clear homology to MCTS1-DENR, a validated mammalian re-initiation factor; the *in vitro* ability shared by both factors to act on 40S subunits and deliver Met-tRNA^Met^_i_ to the empty P-site ([14]; using an experimental model where 40S is in complex with HCV IRES-like mRNA); an *in vitro* role of eIF2D for another viral element-dependent translational shuttling phenomenon that resembles re-initiation [38]; the shared *in vitro* activity to promote the release of deacylated tRNA and mRNA from recycled 40S subunits [14]; the high structural similarities of the proteins in reconstituted complexes together with the 40S ribosome [26, 33, 39]; and, finally, *in vivo* evidence in yeast that the orthologous proteins have shared functions which, however, do not lie in uORF re-initiation but in post-termination tRNA removal and 40S ribosome recycling [19] – events from which re-initiation can be envisaged to have subsequently evolved in higher eukaryotes. Finally, two recent studies have observed that eIF2D loss-of-function affected the expression of ATF4 in the integrated stress response in *Drosophila* [22] and of reporter constructs that carried the *Atf4* 5′ UTR (or several other 5′ UTRs containing uORFs) in HeLa cells [21]. These effects were exacerbated by simultaneous DENR loss-of-function, which was interpreted as partially overlapping re-initiation activities of eIF2D and MCTS1-DENR. However, whether the eIF2D-dependent effects indeed relied on the mRNAs’ uORFs, or whether they could be mechanistically distinct, was not specifically addressed. Notably, evidence has been accumulating that eIF2D plays roles separate from MCTS1-DENR [20, 28, 40, 41], and the data presented in our study supports this idea as well.

Our work, based on the transcriptome-wide analysis of how RNA abundances and translation are altered upon eIF2D depletion, provides evidence that eIF2D is an important regulator of cellular gene expression, as judged from the massive reprogramming at the transcriptome level with hundreds of mRNAs up- and down-regulated and a cell cycle phenotype. However, minor changes occur at the translational level and, moreover, the affected mRNAs are not enriched for uORFs, which points to a main cellular role that does not involve re-initiation and therefore does not overlap with that of MCTS1-DENR. The specific analysis of *Atf4* mRNA in our datasets also supports this model, as we observe increased footprint abundance on the *Atf4* uORF1 stop codon in *Denr* shRNA-treated cells (in line with ribosomal stalling when deacylated tRNA removal is impaired [21]), but not so in *Eif2d* shRNA-treated cells whose profiles are identical to control cells (**Supplementary Figure S6**). Still, since our *in vitro* experiments also reveal that high concentrations of eIF2D can partially compensate for MCTS1-DENR loss, it is quite likely that eIF2D has retained some capacity to act as a re-initiation factor. Whether this activity comes to bear under certain conditions or in specific cell types *in vivo* will require careful further investigations. Probably the most revealing experiment to elucidate the biological function and mechanism by which eIF2D acts, and to distinguish direct from indirect targets in our profiling analyses, would be to identify the mRNAs and sequence elements that eIF2D physically interacts with *in vivo*, for example through selective 40S footprinting [13]. Our efforts in this direction have, unfortunately, not yet been successful.

That MCTS2 can act as a *bona fide*, functionally active DENR interaction partner that is expressed to significant levels in the NIH/3T3 cells used for our study, is an important finding as it potentially explains some of the enigmatic published discrepancies between *Mcts1* vs. *Denr* knockout phenotypes. Thus, neuronal migration has been reported to be more strongly perturbed upon DENR loss-of-function than MCTS1 loss-of-function [29]. Moreover, according to the International Mouse Phenotyping Consortium (IMPC; https://www.mousephenotype.org/), *Denr* knockout leads to preweaning lethality with complete penetrance, yet *Mcts1* knockouts are viable, with relatively mild reported phenotypes (mainly affecting eye morphology in early adult animals). The consortium did not phenotype knockouts of *Mcts2*, likely due to its pseudogene assignment. The high *Mcts2* RNA and footprint expression observed in our study could be a particularity of NIH/3T3 cells. Of note, our pilot experiments in mouse embryonic stem cells (mESCs) have recovered MCTS2 as an *in vivo* interactor of DENR as well (data not shown), and re-analysis of our previous Ribo-seq data from mouse liver [25] and kidney [42] indicates that MCTS2 is co-expressed alongside MCTS1 in these organs *in vivo*, with relative proportions of MCTS1 vs. MCTS2 biosynthesis different across these organs (**Supplementary Figure S7**). Therefore, we propose that MCTS2 is a broadly expressed paralog. In the future, it will be particularly interesting to explore whether MCTS1-DENR and MCTS2-DENR heterodimers have redundant functions or whether the identity of the MCTS paralog has functional consequences for the activity of the heterodimer. Finally, while Bohlen et al. recently reported that *Mcts2* expression is undetectable in various cells of the myeloid lineage (human whole blood cells, THP-1 cells, T-cell blasts) [32], our findings from mouse warrant a more thorough analysis across human tissue and cell types.

## Material and methods

### Cloning

To generate *in vivo* dual luciferase reporter plasmids, we amplified the 5′ UTRs and first 5 or 10 (depending on construct) CDS codons (in frame with firefly luciferase CDS) of the selected transcripts by PCR from mouse cDNA or genomic DNA and cloned them into the BamHI restriction site of pLV1 dual luciferase reporter plasmid with mutated firefly CDS start, essentially as previously described [17]. Sequences of the primers used for the PCR are reported in Table 1. All constructs were validated by Sanger sequencing.

*In vitro* translation reporters were generated by PCR amplification of *Asb8* or *Klhdc8a* 5′ UTRs together with the 6 first CDS codons (in frame with the firefly luciferase CDS, lacking additional ATG start codon), from mouse cDNA, genomic DNA, or the corresponding pLV1 *in vivo* reporter constructs. The PCR products were then cloned into the BamHI restriction site of luciferase T7 control DNA plasmid (Cat. No. L4821, Promega). Modifications of uORFs or other mutations were performed using the Q5 site-directed mutagenesis kit (Cat. No. E0554S, New England Biolabs) and validated by Sanger sequencing. Sequences of primers for PCRs and mutagenesis are reported in Table 2.

Plasmids for MCTS1, DENR and eIF2D protein expression have been reported [33] and cloning of human *Mcts2* cDNA for recombinant expression was analogous to that of *Mcts1*; briefly, a fragment encoding the first 154 amino acids of MCTS2 (i.e., the part carrying substitutions with regard to MCTS1) was PCR-amplified from HeLa genomic DNA with primers reported in Table 3 and cloned into the XhoI and EcoRV restriction sites of the pLIC C-terminally 6×His-TEV-tagged MCTS1-expression vector [33]. Correct sequence was validated by Sanger sequencing.

For endogenous tagging of DENR, a gRNA (GTCATCCACCTAGAAGACACAGG) targeting a downstream sequence of the *Denr* stop codon was inserted into vectors pCas9-2A-mCherry [43] and pRRE200 [44] using oligos described in Table 4.5 µM of oligos were annealed in annealing buffer (100 µM Tris-HCl pH 7.5, 500 mM NaCl and 10 mM EDTA) by cooling from 98 °C to 20 °C with a ramp of 0.02 °C per second. Annealed oligos were diluted 10 times then ligated into pCas9-2A-mCherry BsaI restriction site or pRRE200 AatII and SacI restriction sites. Colony PCRs using the reverse gRNA oligo and a pCas9-2A-mCherry or pRRE200 forward primer was carried out to check for correct insertion of the annealed oligos and the cloned plasmids validated by Sanger sequencing.

A pBluescript II SK(-) vector was ordered as a synthetic clone to contain the knock-in cassette 3×FLAG-NeonGreen-dTAG. *Denr* 5′ and 3′ homology arms were amplified from mouse genomic DNA by PCR using primers reported in Table 4 and ligated in the pBluescript II SK(-) first digested with SpeI and BamHI for the 5′ homology arm then with NdeI and KpnI for the 3′ homology arm. Positive clones were identified by colony PCR using PCR homology arm primers and by Sanger sequencing.

**Table 1.**
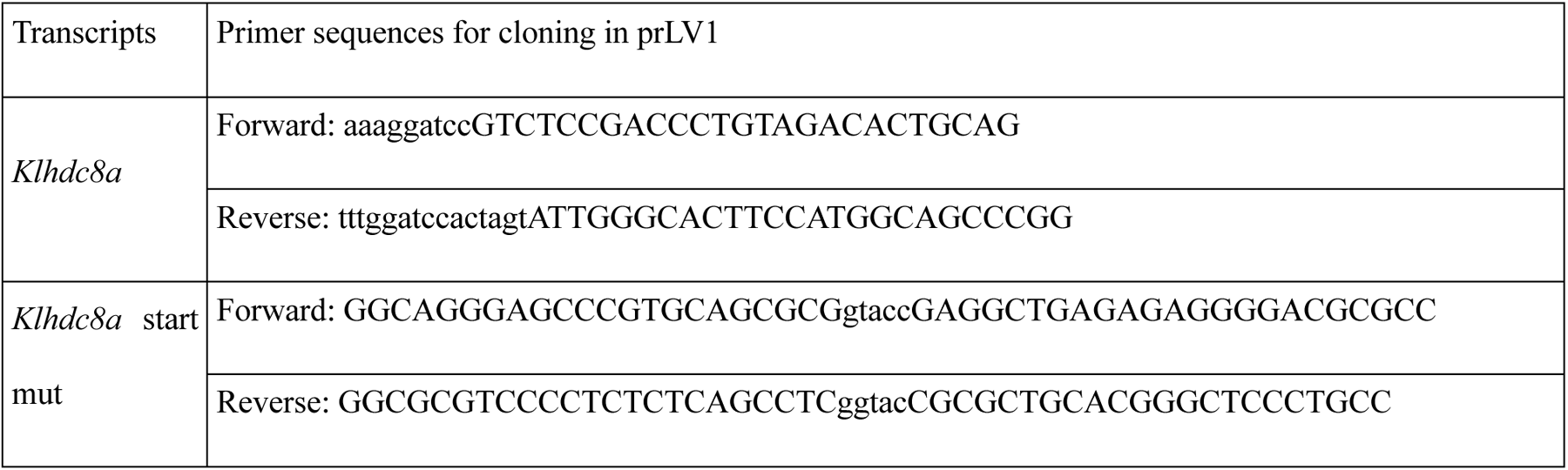
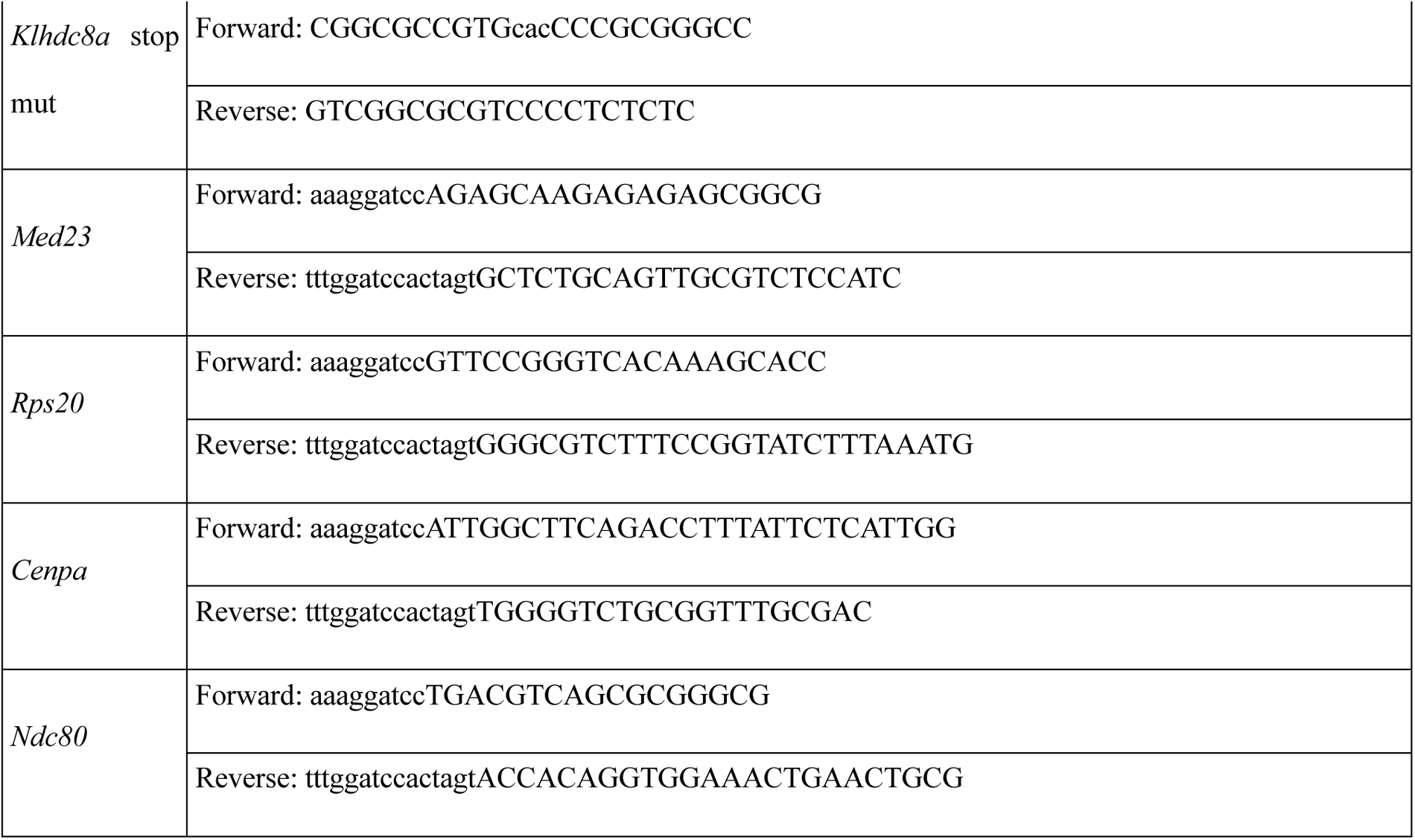
Sequences of oligonucleotide primers used for the *in vivo* reporters.

**Table 2.**
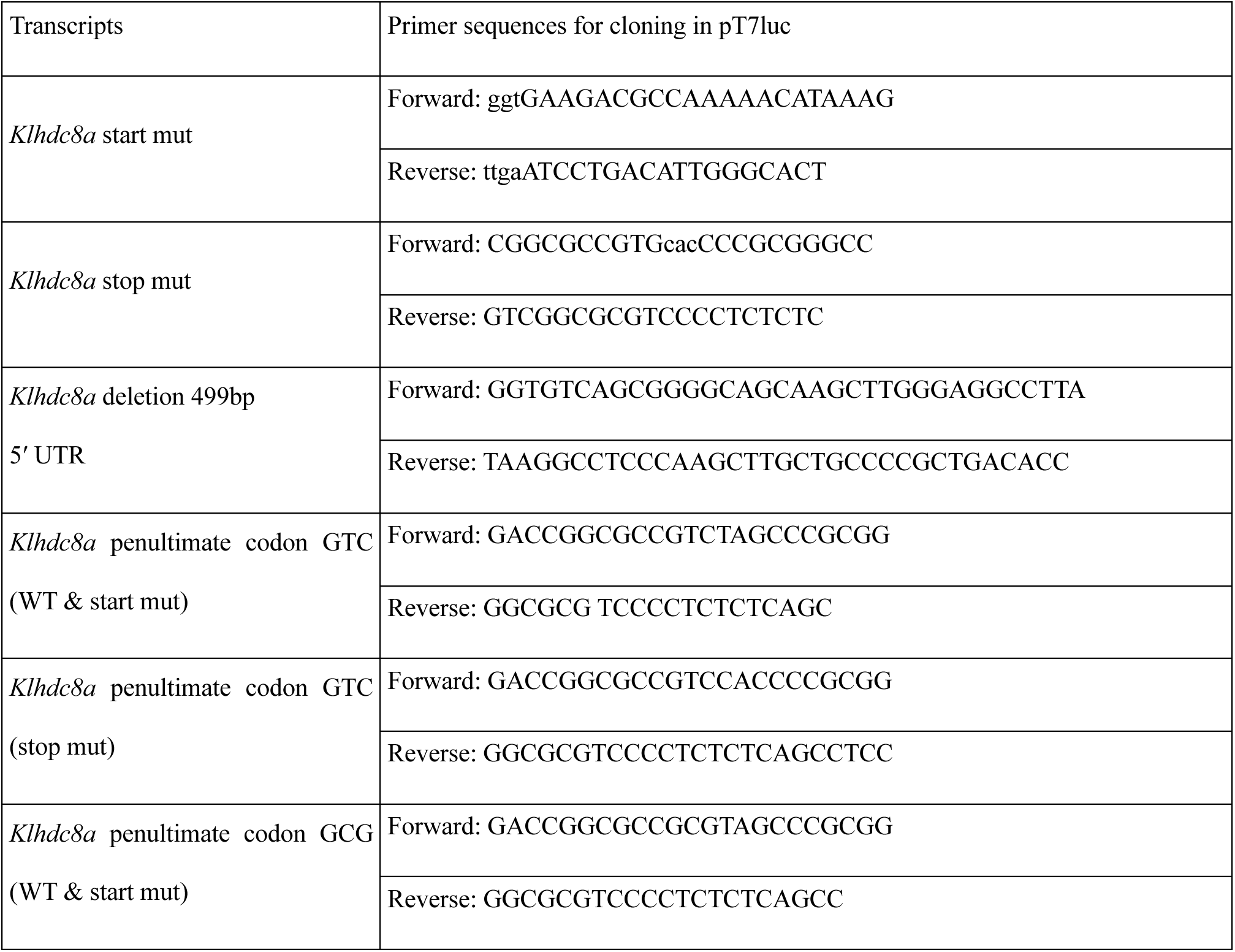
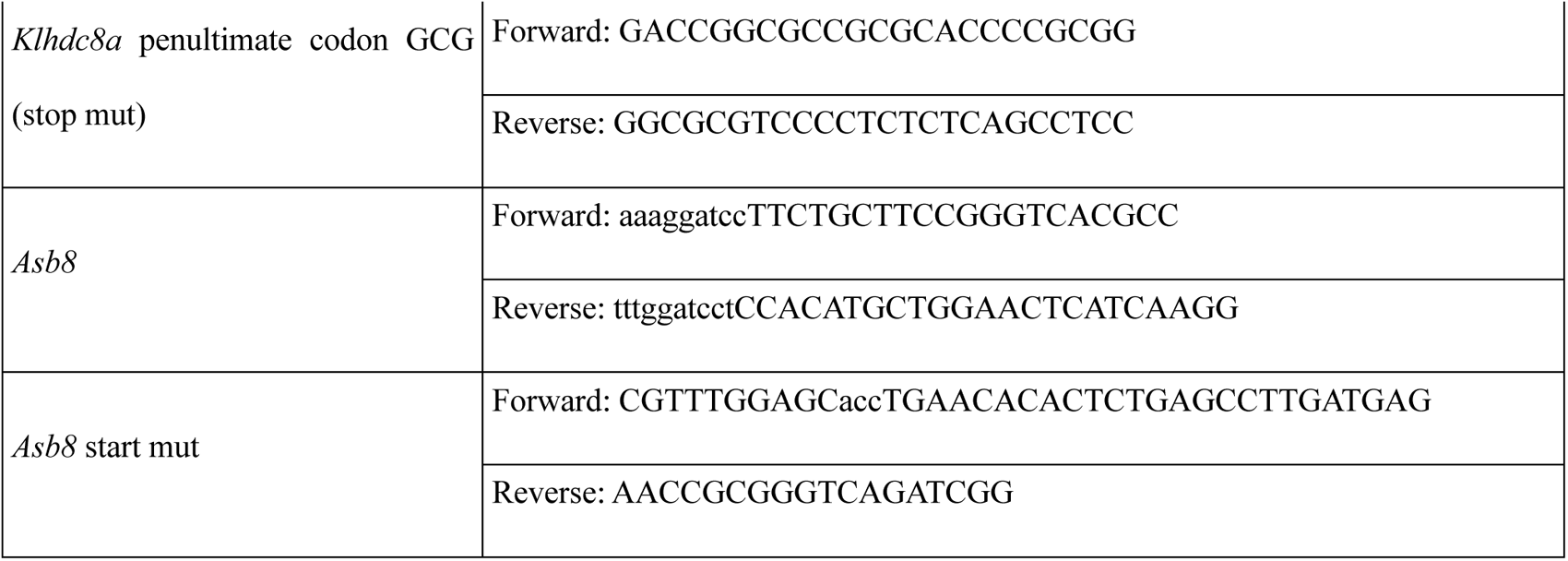
Sequences of oligonucleotide primers used for the *in vitro* reporters.

**Table 3.**
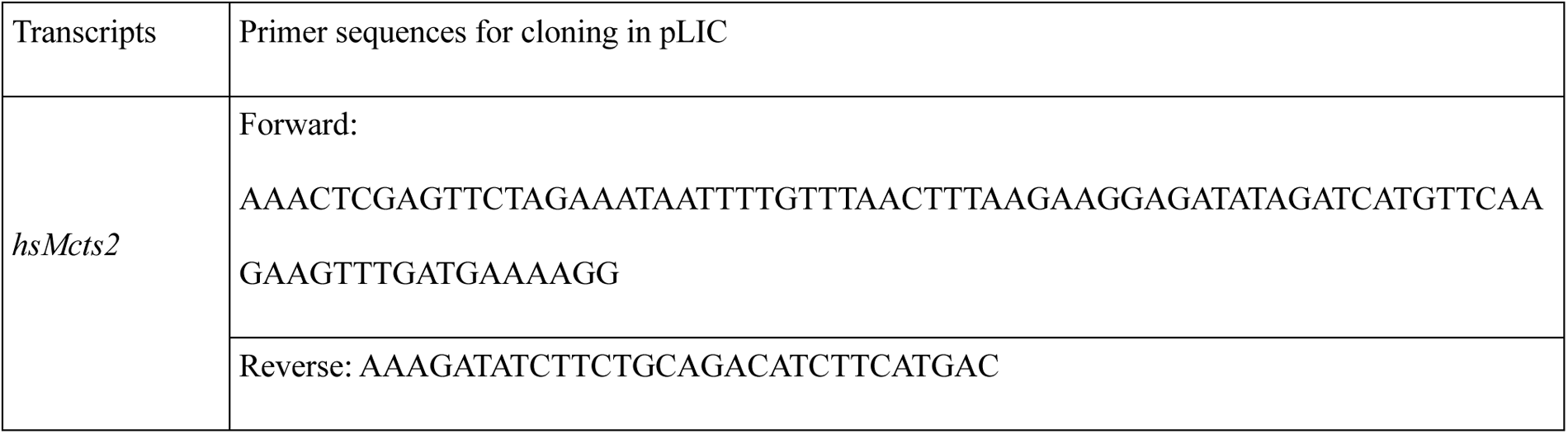
Sequences of oligonucleotide primers used for cloning the pLIC C-terminally 6x-His-TEV-tagged *Mcts2* vector.

**Table 4.**
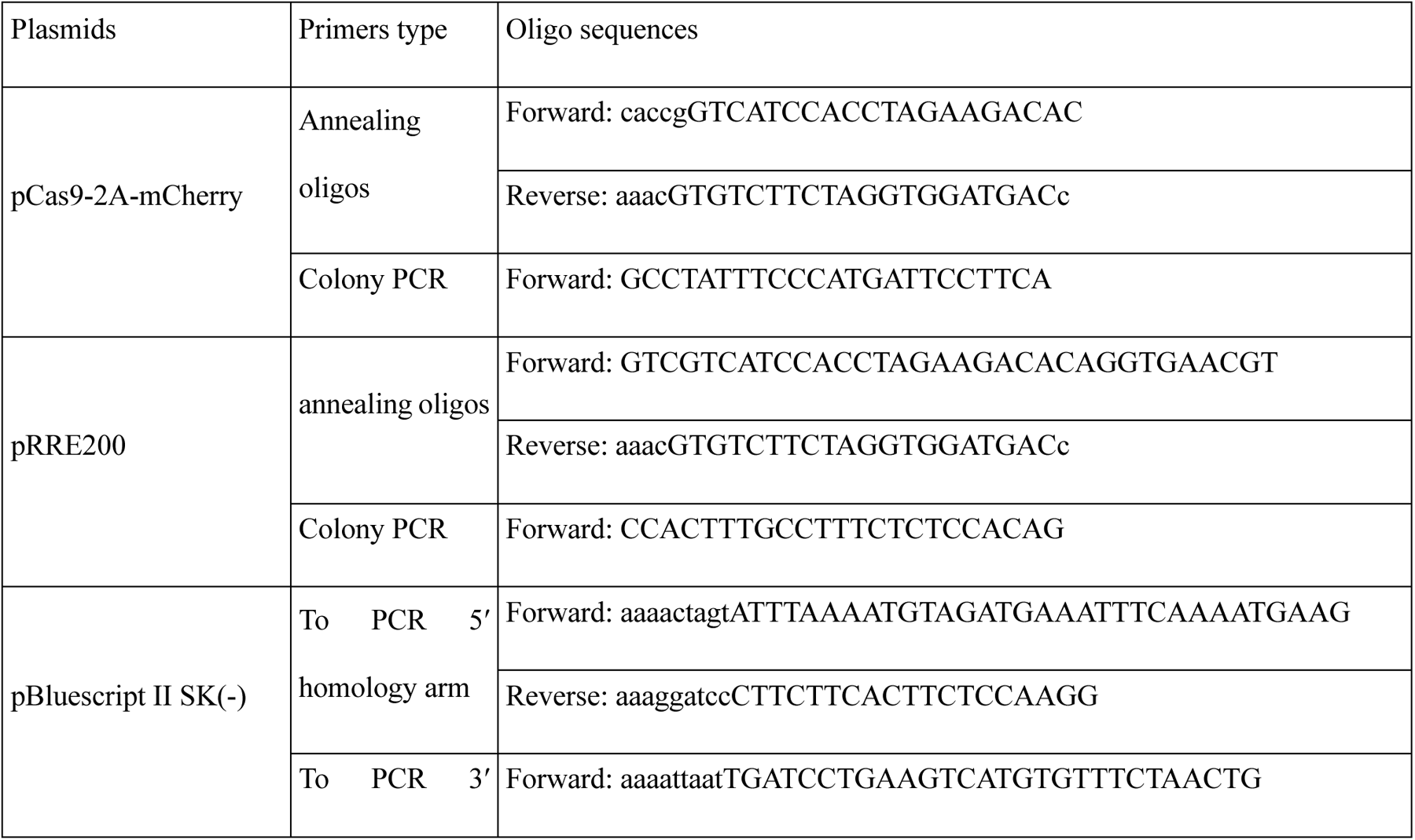
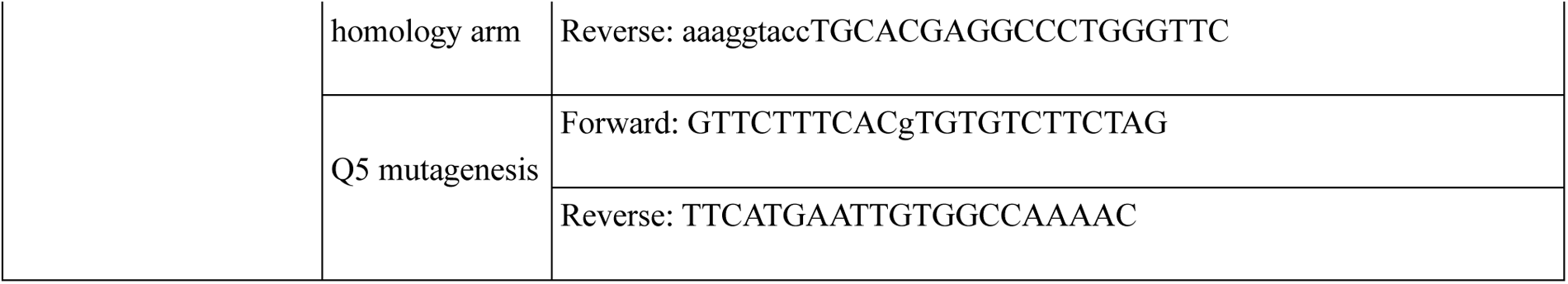
Sequences of oligonucleotides used for the cloning of *Denr* gRNA and homology arms into the pCas9-2A-mCherry, pRRE200 and pBluescript II SK(-).

### Cell culture and generation of HeLa knockout cells

NIH/3T3, HEK293FT and HeLa cells were cultured under standard conditions (DMEM; 10 % FCS, 1 % penicillin/streptomycin, all from Invitrogen; 37 °C; 5 % CO_2_). The HeLa *Denr* knockout (KO) cells have been published [21]. For *Eif2d* KOs, wild-type HeLa cells were seeded in a 6-well plate (200,000 cells per well) in DMEM with 10 % FBS and 1 % Pen/Strep. After 24 h the cells were transfected with plasmid pX459 v2.0 into which an sgRNA sequence targeting *Eif2d* was cloned (sequence: CACCGCTGTGGACTGGAAACACCCG) using Lipofectamine 2000 following the manufacturer’s instructions. 24 h later, lipofectamine was removed and cells were kept in DMEM for recovery for 2 days. On the third day, puromycin was added to a final concentration of 1.5 µg/ml. After another 3 days, puromycin was removed and fresh medium was added. After 2 days, cells were seeded into a 96-well plate at limiting dilution, so that only one cell per well was seeded. Wells were checked under the microscope for single cell colonies. Upon reaching confluency wells with only one colony were reseeded into 6-well plate and kept for growing. Then immunoblots on these monoclonal lines were performed to identify the ones with loss of eIF2D. The knockout was verified at the genomic level via PCR using the following primers: 5′-GGGGAGGGCTCGGGTATGAC-3′ and 5′-AAGGCAGGGGCTGTCTCATATC-3′ to amplify the genomic region. This revealed premature stop codons in all alleles. For generating *Eif2d / Denr* double knockout cells, the same procedure was performed to introduce the *Eif2d* knockout into the previously published *Denr*-knockout HeLa line [21].

### Lentiviral transduction

Lentiviral shRNA plasmids targeting *Eif2d* were purchased from Sigma (Cat. No. TRCN0000119842 for non-functional (nf) shRNA, TRCN0000119844 for shRNA1 and TRCN0000119845 for shRNA2). The *Denr* (“*Denr* shRNA2”) and Scramble shRNA plasmids have been described previously [17]. Production of lentiviral particles in HEK293FT cells using envelope pMD2.G and packaging psPAX2 plasmids and viral transduction of NIH/3T3 cells were performed following published protocols [17, 45], with puromycin selection at 5 μg/ml for 4 days for shRNA transduced cells.

### Western Blot

Total protein extracts from NIH/3T3 and HeLa cells were prepared according to the NUN method [46]. For analysis of phosphorylated proteins, PhosSTOP™ (Cat. No. 4906845001, Roche) was included in the NUN buffer. Antibodies used were rabbit anti-KLHDC8A 1:5000 (ab235419, Abcam), rabbit anti-VINCULIN 1:5000 (ab129002, Abcam) mouse anti-U2AF65 1:5000 (U4758, Sigma), rabbit anti-EIF2D 1:750 (12840-1-AP, Proteintech), rabbit anti-DENR 1:5000 (ab108221, Abcam), rabbit anti-phospho-eIF2alpha (Ser51) 1:1000 (#9721S, Cell Signaling), mouse anti-FLAG 1:10000 (M2F3165, Sigma), rabbit anti-RPL23 (ab112587, Abcam), mouse anti-beta-Tubulin 1:5000 (T5201, Sigma) and secondaries goat anti-guinea pig-HRP 1:10000 (SA00001-12, Proteintech), anti-rabbit-HRP (W4011, Promega) and anti-mouse-HRP 1:10000 (S3721, Promega). The guinea-pig anti-MCTS1 antibodies (used at 1:500) have been described (Schleich et al. 2014). Blots were visualized with the Fusion Fx7 system.

### Translation-competent extract preparation

The overall protocol for extract preparation was an adaptation of that published by Gurzeler et al., 2022 [23]. Briefly, confluent 15-cm dishes of HeLa cells were washed once with PBS at 37 °C, then trypsinized with 0.05 % Trypsin-EDTA (Cat. No. 25300054, Gibco™). Harvested cells were collected by centrifugation at 500 × *g* for 5 min at 4 °C. The cell pellet was washed twice with ice-cold PBS and centrifugation at 200 × *g* for 5 min at 4 °C. The cell pellet was resuspended in 1:1 lysis buffer (10 mM HEPES pH 7.3, 10 mM potassium acetate, 500 μM MgCl_2_, 5 mM DTT and 1× cOmplete™ EDTA-free Protease Inhibitor Cocktail (Cat. No. 4693132001, Roche)). Cells were lysed by dual centrifugation at 500 RPM for 4 min at −5 °C using a ZentriMix 380 R centrifuge (Hettich AG) with a 3206 rotor and 3209 adapters. The resulting lysate was then centrifuged in a table-top centrifuge at 13,000 × *g* at 4 °C for 10 min and the supernatant was aliquoted and snap-frozen in liquid nitrogen; storage at −80 °C for up to 6 months.

### Expression and purification of recombinant proteins

Recombinant EIF2D, MCTS1 and DENR protein purification has been described [33]. Recombinant MCTS2 was expressed and purified identically to MCTS1. GADD34Δ1-240 expression plasmid has been described [23]; for protein preparation, bacterial strain BL-21 (DE-3) (pLysS) was transformed with the plasmid, grown in 1000 ml Luria broth at 37 °C until the OD600 reached 0.4–0.6. After induction (1 mM IPTG) bacteria were cultured for another 4 h and cells were harvested. Expression of recombinant GADD34Δ1-240 was validated on Coomassie gel. The cell pellet was resuspended in 50 ml lysis/wash buffer (50 mM Tris-HCl pH 7.5, 300 mM NaCl, 5 % glycerol, 25 mM imidazole), lysed by sonication with 240 × 1 s pulses at 40 % amplitude on ice and centrifuged at 18,000 × *g* for 20 min at 4 °C. The supernatant was mixed with 1 ml HisPur Ni-NTA washed resin slurry (Cat. No. 88222, Thermo Fisher Scientific) for at least 2 h at 4 °C under continuous rotation. Unbound proteins and background were removed by washing the resin 2× with 50 ml of lysis/wash buffer, then bound proteins were eluted 2× with 3 ml elution buffer (lysis/wash buffer supplemented with 400 mM imidazole). The proteins were dialyzed in protein reconstitution buffer (30 mM NaCl, 5 mM HEPES pH 7.3) overnight using a Slide-A-Lyzer Dialysis Cassette (MWCO 10K) (Cat. No. 88400TS, Thermo Fisher Scientific) at 4 °C under agitation. Protein concentration was quantified with a nanodrop device.

### *In vitro* transcription

Reporter DNA templates were PCR-amplified before transcription for 2 h using the MEGAscript® T7 Transcription Kit protocol (AM1334, Invitrogen) (forward primer: GCGCGTTGGCCGATTCATTAATGC; reverse primer: T_100_GGGAGCTCGCCCCCTCGGAG). Capping was carried out co-transcriptionally using the CleanCap Reagent AG (3′ Ome) (N-7413, TriLink) and the Yeast Inorganic Pyrophosphatase (Cat. No. M2403S, New England BioLabs) following the CleanCap protocol. The capped RNAs were treated for 15 min at 37 °C with 0.14 U/μl Turbo DNase (AM2238, Invitrogen), then purified using the Monarch RNA Cleanup Kit (T2040L, New England Biolabs). Quality and size of the produced RNAs were assessed using an Agilent Fragment Analyzer.

### *In vitro* translation assay

6.5 µl of translation-competent extracts were freshly supplemented with 0.4 mM amino acids (L4461, Promega), 15 mM HEPES pH 7.3, 6 mM Creatine Phosphate, 102 ng/ml Creatine kinase, 28 mM potassium acetate, 1 mM MgCl_2_, 24 mM KCl, 1 U/µl RNasin® Plus Ribonuclease Inhibitor (N2611, Promega) and 2 fmol/µl of Renilla luciferase reporter in a final volume of 12.5 µl prior to each *in vitro* translation assay. Unless specified, 16 ng/µl of recombinant GADD34Δ1-240 was included to the supplemented extracts. 30 fmol/µl of firefly luciferase reporter were used per reaction. The *in vitro* translation assays were carried out at 37 °C for 50 min. Luminescence activity was quantified using the Dual-Glo® Luciferase Assay System (E2940, Promega) and a Tecan Safire2™ plate reader.

### *In vivo* dual luciferase assay

NIH/3T3 cells were first stably transduced with the lentiviral dual luciferase reporters containing the 5′ UTRs to be assayed (upstream of firefly luciferase) and Renilla luciferase (internal normalisation), bidirectionally driven by the *Pgk1* promoter. After establishing the reporter-expressing cell population and several passages to ensure stable expression, cells were transduced with the shRNA lentiviruses and selected on puromycin for 4 days before lysis in 1× passive lysis buffer (E1941, Promega) and dual luciferase readout using the Dual-Glo® Luciferase Assay System (E2940, Promega) and a Tecan Safire2™ plate reader.

### Ribosome profiling and library preparation

One 15-centimeter dish of confluent NIH/3T3 cells was used per replicate. Ribosome profiling and parallel RNA-seq were performed in triplicate for *Denr* shRNA2, *Eif2d* shRNA1, *Eif2d* shRNA2, non-functional (nf) *Eif2d* shRNA and Scr shRNA transduced cells, closely following our previously reported protocol [17] with minor modifications as described below. Cells were washed in cold PBS (without cycloheximide), harvested by scraping down in small volume of PBS (without cycloheximide), pelleted by brief centrifugation, then flash frozen. Cell lysates were prepared as described (with cycloheximide) [17]. For the generation of ribosome-protected fragments, cell lysates were treated with 5 units RNaseI (ART-Seq, Epicenter/Illumina) per OD260, and monosomes were purified on MicroSpin S-400 columns (GE Healthcare). For both Ribo-and RNA-seq, 2 µg of RNA was used for rRNA depletion following the RiboCop rRNA Depletion Kit V1.2 (Cat. No. 037.24, Lexogen). Finally, libraries were amplified using 10-14 PCR cycles, and sequenced (HiSeq2500 platform).

### Sequencing pre-processing, alignment, quantification

Sequencing reads were processed as described previously [47]. Briefly, sequencing adapters were trimmed using Cutadapt [48] and the reads were size filtered by a custom Python script (26-35 nt for RPF and 21-60 nt for RNA). The reads were then aligned subsequently to mouse rRNA, human rRNA, mouse tRNA and mouse cDNA from Ensembl mouse database release 100 using Bowtie2 [49]. In order to estimate expressed isoforms, the RNA-seq reads were mapped in parallel to the mouse genome using STARmapper version 2.5.3a [50] and processed with StringTie [51] to measure FPKM for each transcript. A custom Python script classified the transcripts into single expressed isoform or multiple transcript isoforms. The mapped reads were counted on 5′ UTR or CDS of the protein coding genes using a custom Python script. The location of the putative A-site of RPFs were estimated as 5′ position +16 for reads shorter than or equal to 31 nt and +17 for reads longer than 31 nt. Transcripts for which there were less than 10 counts in total of all samples were filtered out of the further analysis. Read counts were normalized using the upper quartile normalization method [52] from edgeR [52] then transformed to transcripts per million (TPM) to normalize for transcript length. Translation efficiencies were calculated as the log_2_-transformed ratio of normalized CDS footprint counts to normalized CDS RNA counts and averaged over replicates. All analyses were carried out on R version 4.2.1.

### Differential translation efficiency analysis

Ribosome profiling sequencing data were analysed using the deltaTE method [53] in order to assess significant changes in translation efficiency and RNA abundance. Significance threshold was set to FDR < 0.1.

### uORF annotation and analyses

For uORF annotation, transcripts were used for which reads could be unambiguously assigned to the 5′ UTR, i.e. transcripts that are the only protein-coding isoform expressed (single expressed isoform) (N = 6841). In order to include as many transcripts as possible for the annotations, genes with multiple protein coding isoforms were also considered if all expressed isoforms had the same CDS start, in which case the transcript with the longest 5′ UTR was selected (N = 2425). For the selected transcripts, uORFs were annotated and considered as translated with the following criteria: (i) they started with AUG, CUG, GUG or UUG, (ii) they had an in-frame stop codon within the 5′ UTR, and (iii) they had a coverage of at least 33 %. When several potential uORFs were overlapping, the one with the highest coverage (read count/uORF length) was considered.

### Cell cycle monitoring

The cell proliferation assay was performed using the Click-iT® EdU Flow Cytometry Assay Kits (C10424, Invitrogen) following the vendor’s protocol with some modifications. Briefly, NIH/3T3 cells transduced with scramble shRNA, nf *Eif2d* shRNA, *Denr* shRNA and *Eif2d* shRNA1 and shRNA2 were seeded into 6-cm plates under puromycin selection. After 4 days, the cells were treated for 1 h with 10 µM of EdU, washed with PBS, trypsinized and collected into FACS tubes. The cells were then centrifuged at 1200 × *g* for 5 min and resuspended in PBS. The cells were counted, and all conditions were adjusted to the same number of cells. After washing with 3 ml of 1 % BSA in PBS, the cells were resuspended in 100 µl fixative buffer and incubated overnight at 4 °C. Another wash with 1 % BSA in PBS followed by a 15 min incubation in 100 µl of 1×-saponin-based permeabilization and wash reagent. 200 µl of Click-iT reaction cocktail were added to each tube and the cells were incubated for 30min at room temperature. After a wash in 3 ml of 1 % BSA in PBS, the cells were resuspended in 100 µl of 1×-saponin-based permeabilization and wash reagent complemented with 5µl of 20 mg/ml RNaseA and 2 µl of 1 mg/ml propidium iodide and incubated for 30 min at room temperature. Fluorescence was measured using Accuri C6 FACS machine.

### Generation of endogenously tagged cell lines

C-terminal tagging of endogenous DENR with 3×FLAG-NeonGreen-dTAG in NIH/3T3 cells was carried out using CRISPR-Cas9 homology-directed repair. 10^5^ NIH/3T3 cells were seeded in a 6-well plate 24h prior to transfection. 0.9 µg of the pBLU donor plasmid (containing 400 bp 5′ and 3′ homology arms directly upstream and downstream of *Denr* stop codon) as well as 0.5 µg of the pCas9-gRNA and 0.1 µg of the control pRRE reporter were transfected into NIH/3T3 using Lipofectamine™ 3000 (L3000008, Invitrogen). Medium was changed after 24 h and cells were FACS sorted based on mCherry and EGPF expression in a 10 cm dish 48 h post-transfection. Cells were counted and diluted to have 1 cell per well of a 96-well plate. When cell clones became visible, they were preselected based on NeonGreen expression, then split into a 96-well plate for genotyping and a 12-well plate for maintenance. For genomic DNA extraction, cells were washed with PBS then lysed in 20 µl of DNA lysis buffer (10 mM Tris-HCl pH 8, 0.5 mM EDTA, 0.5 % Triton X-100, 0.5 mg/ml proteinase K). After incubation for 1 h at 55 °C, cooling on ice and further incubation for 10 min at 95 °C to inactivate proteinase K, this prep was used to check the insertion of the cassette into the *Denr* locus by PCR using primers upstream of the 5′ and downstream of the 3′ homology arms.

### Co-immunoprecipitation

Triplicate 15 cm plates of 80 % confluent NIH/3T3 DENR-3×FLAG-NeonGreen-dTAG cells were treated with 500 nM dTAG-13 or control-treated for 5 h prior to harvesting. After washing with PBS, cells were scraped in 1 ml PBS (4 °C). Cells were collected by centrifugation at 500 × *g* for 5min at 4 °C and lysed in 400 µl IP buffer (50 mM Tris-HCl pH 7.5, 120 mM NaCl, 1 % NP-40, 0.5 mM DTT, 10 mM MgCl_2_, 1× PhosSTOP™ (4906837001, Roche), 1× cOmplete™ EDTA-free Protease Inhibitor Cocktail (Cat. No. 4693132001, Roche), 0.04 U/µl RNasin® Plus Ribonuclease Inhibitor (CN2611, Promega)) for 30 min on ice with mixing by pipetting every 10 min. Lysed cells were centrifuged for 7 min at 5000 × *g* at 4 °C and the supernatant was collected. Proteins were quantified using Pierce™ BCA Protein Assay kit (Cat. No. 23225, Thermo Fisher Scientific) and 500 µl of 2 mg/ml proteins were incubated with washed anti-FLAG magnetic beads (Cat. No. M8823, Millipore) at 4 °C for 3 h on rotating wheel. The flow-through was kept for Western Blot and beads were washed twice in wash buffer (20 mM Tris-HCl pH 7.5, 100 mM NaCl, 0.5 mM DTT, 10 mM MgCl_2_, 1× cOmplete™ EDTA-free Protease Inhibitor Cocktail (Cat. No. 4693132001, Roche), 0.04 U/µl RNasin® Plus Ribonuclease Inhibitor (N2611, Promega)) with change of tubes between the 1^st^ and the 2^nd^ wash. Proteins were eluted in 80 µl elution buffer (NH_4_OH pH 11-12) and incubated for 15 min at 4 °C. Elution step was repeated twice and eluates were pooled, frozen and dried in a SpeedVac system.

### Mass-spectrometry and data analysis

Samples were digested following a modified version of the iST method (named miST method). Briefly, dried material was redissolved in 50 µl miST lysis buffer (1 % Sodium deoxycholate, 100 mM Tris pH 8.6, 10 mM DTT), heated 5 min at 95 °C and diluted 1:1 (v:v) with water. Reduced disulfides were alkylated by adding ¼ vol. of 16 0mM chloroacetamide (32 mM final) and incubating for 45 min at RT in the dark. Samples were adjusted to 3 mM EDTA and digested with 1.0 µg Trypsin/LysC mix (Promega #V5073) for 1 h at 37 °C, followed by a second 1 h digestion with an additional 0.5 µg of proteases. To remove sodium deoxycholate, two sample volumes of isopropanol containing 1 % TFA were added to the digests, and the samples were desalted on a strong cation exchange (SCX) plate (Oasis MCX; Waters Corp., Milford, MA) by centrifugation. After washing with isopropanol/1 % TFA, peptides were eluted in 200 µl of 80 % MeCN, 19 % water, 1 % (v/v) ammonia, and dried by centrifugal evaporation. Tryptic peptide mixtures were injected on an Ultimate RSLC 3000 nanoHPLC system interfaced via a nanospray Flex source to a high resolution Orbitrap Exploris 480 mass spectrometer (Thermo Fisher, Bremen, Germany). Peptides were loaded onto a trapping microcolumn Acclaim PepMap100 C18 (20 mm × 100 μm ID, 5 μm, Dionex) before separation on a C18 custom packed column (75 μm ID × 45 cm, 1.8 μm particles, Reprosil Pur, Dr. Maisch), using a gradient from 4 % to 90 % acetonitrile in 0.1 % formic acid for peptide separation (total time: 140 min). Full MS survey scans were performed at 120,000 resolution. A data-dependent acquisition method controlled by Xcalibur software (Thermo Fisher Scientific) was used that optimised the number of precursors selected (“top speed”) of charge 2+ to 5+ while maintaining a fixed scan cycle of 2 s. Peptides were fragmented by higher energy collision dissociation (HCD) with a normalized energy of 30 % at 15,000 resolution. The window for precursor isolation was of 1.6 *m/z* units around the precursor and selected fragments were excluded for 60s from further analysis. Data files were analysed with MaxQuant 1.6.14.0 incorporating the Andromeda search engine Cysteine carbamidomethylation was selected as fixed modification while methionine oxidation and protein N-terminal acetylation were specified as variable modifications. The sequence databases used for searching were the mouse *(Mus musculus)* reference proteome based on the UniProt database (www.uniprot.org, version of April 6^th^, 2021, containing 55,341 sequences RefProt _Mus_musculus_20210604.fasta) supplemented with the sequences of common contaminants, and a database of mammalian immunoglobulin sequences (https://www.imgt.org version of 20th Nov. 2018, 2243 entries). Mass tolerance was 4.5 ppm on precursors (after recalibration) and 20 ppm on MS/MS fragments. Both peptide and protein identifications were filtered at 1 % FDR relative to hits against a decoy database built by reversing protein sequences. All subsequent analyses were done with the Perseus software package (version 1.6.15.0). Contaminant proteins were removed, and intensity iBAQ values were log_2_-transformed. After assignment to groups, only proteins quantified in at least 3 samples of one group were kept. After missing values imputation (based on normal distribution using Perseus default parameters), *t*-tests were carried out among all conditions, with permutation-based FDR correction for multiple testing (Q-value threshold < 0.05). Log_2_FC of non-treated Denr-3×FLAG-NeonGreen-dTAG vs. dTAG treated Denr-3×FLAG-NeonGreen-dTAG was calculated and p-value was corrected for multiple testing by False discovery rate (significance threshold was set to FDR < 0.05).

## Acknowledgements

We thank the Lausanne Genomic Technologies Facility for library sequencing and the Lausanne Protein Analysis Facility for mass spectrometry-based proteomics. Evan Karousis for help with the initial setup of the *in vitro* translation assay and for the GADD34Δ1-240-expression plasmid, and Michael Taschner for his help with preparation of recombinant GADD34Δ1-240. We thank members of the Gatfield lab for critical comments on the manuscript.

## Author contributions

Conceptualization: D.G., R.M. Methodology: R.M., M.D.M., A.B., T.W. Investigation: R.M., M.D.M., N.G.

Visualization: R.M., N.G., A.B., D.G. Supervision: D.G., N.B., A.A.T. Writing original draft: R.M., D.G.

## Data and script availability

Sequencing data have been deposited at GEO (GSE263991) with reviewer token: *(see cover letter)* and mass spectrometry data at PRIDE (PXD051482), with username: and password:

*(see cover letter)*.

