## Supplementary Figures for "An *in vitro* assay of MCTS1-DENR-dependent re-initiation and ribosome profiling uncover the activity of MCTS2 and distinct function of eIF2D"

A mouse *Klhdc8a* 5' UTR and beginning of CDS (ENSMUST00000046071)

```
  1 GTCTCCGACCCTGTAGACACTGCAGGCCCTGCCCTCGCTCGCCCTGGTCGCCCGGGGCTG   60
 61 ACGCACAGGCCACAGCCTCCGCCTCGGGGCGCGTGGGCTCCTGCCCAGGTGTGGAAGCTG  120
121 CGGGGAGGCGGTGCGGGGAACCCGCGTCCTCCGGCCGGAGCTCGGCTTCTGGCGACCGCT  180
181 CTCCTGCGCGGGTGACGGCGGCGTGCGGGCTGGGAAGGGCAGGGAGCGCTGTCCTCCGCT  240
241 CCGCGGCTGCCAGGCGGACCGTGAAAGCTTCGCGCGGCTGTAGGCGCCGCCCGGGAGCTA  300
301 GTGCTGGGCGAGCGCGGCGCCCACGCGCGGTGGGGTGGGCTCCGAGCTCCGGGGGACCGG  360
361 GCGGGGGCGGGGCGCATCCTGCAAAGTTGACCAAGGGTGAGCTACCGCCGGCGCCCTGCT  420
421 GCCCTAGATTCGCCAGGTTCTTTCCGCGGAGCCAGGCGTGGACGTCGCTGCCCCGCTGAC  480
481 ACCCAAGGGACCCGGAGAAGCCAGCGGGCGTCGGCAGGGAGCCCGTGCAGCGCGAGATGG  540
                                                    uORF -M--
541 AGGCTGAGAGAGGGGACGCGCCGACCGGCGCCGTGTAGCCCGCGGGCCCCGGGGCTGGAC  600
    E--A--E--R--G--D--A--P--T--G--A--V--*-
601 CTTGCCCGGACGCTGTGGGGGCGCTCGAGCAGGGCCCTCCCCCGGGCTGCCATGGAAGTG  660
                                                    CDS -M--E--V-
661 CCCAATGTCAAGGACTTTCAGTGGAAGCGCCTTGCTCCACTGCCTAGCCGCCGGGTCTAT  720
    -P--N--V--K--D--F--Q--W--K--R--L--A--P--L--P--S--R--R--V--Y-
```

B mouse *Asb8* 5' UTR and beginning of CDS (ENSMUST00000143400)

```
  1 AGAATATCATGTCGCTCAGAGGCTCTGACCATGCAGCACCCAGAGAGCACAAGGGGTCTG   60
 61 GGCCAAGAGGGAGGGGAATTTCCTGGAGAAGCACTGAAGATCATGGAAGCAGGTGCAGA  120
121 GGCCCAGGAGCGTGGCTGATCCAGTCCAATTCCAGGCAGTTTGGAGCATGTGAACACACT  180
                                          uORF -M--*-
181 CTGAGCCTTGATGAGTTCCAGCATGTGGTATATTATGCAGAGCATTCAGAGCAAATACTC  240
      CDS -M--S--S--S--M--W--Y--I--M--Q--S--I--Q--S--K--Y--S
```

C

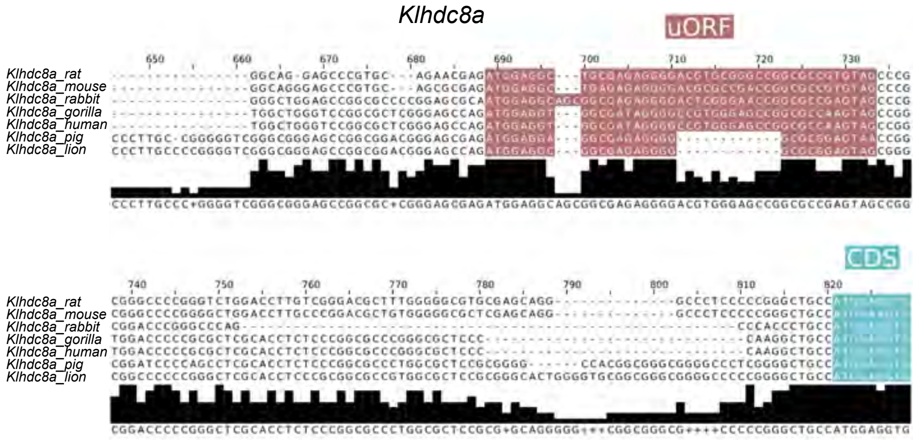


D

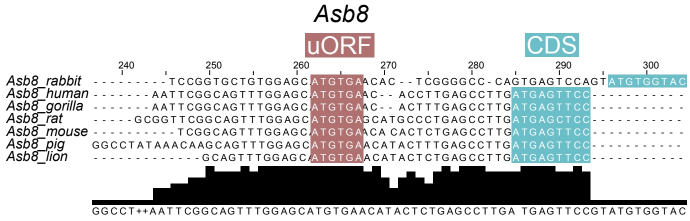

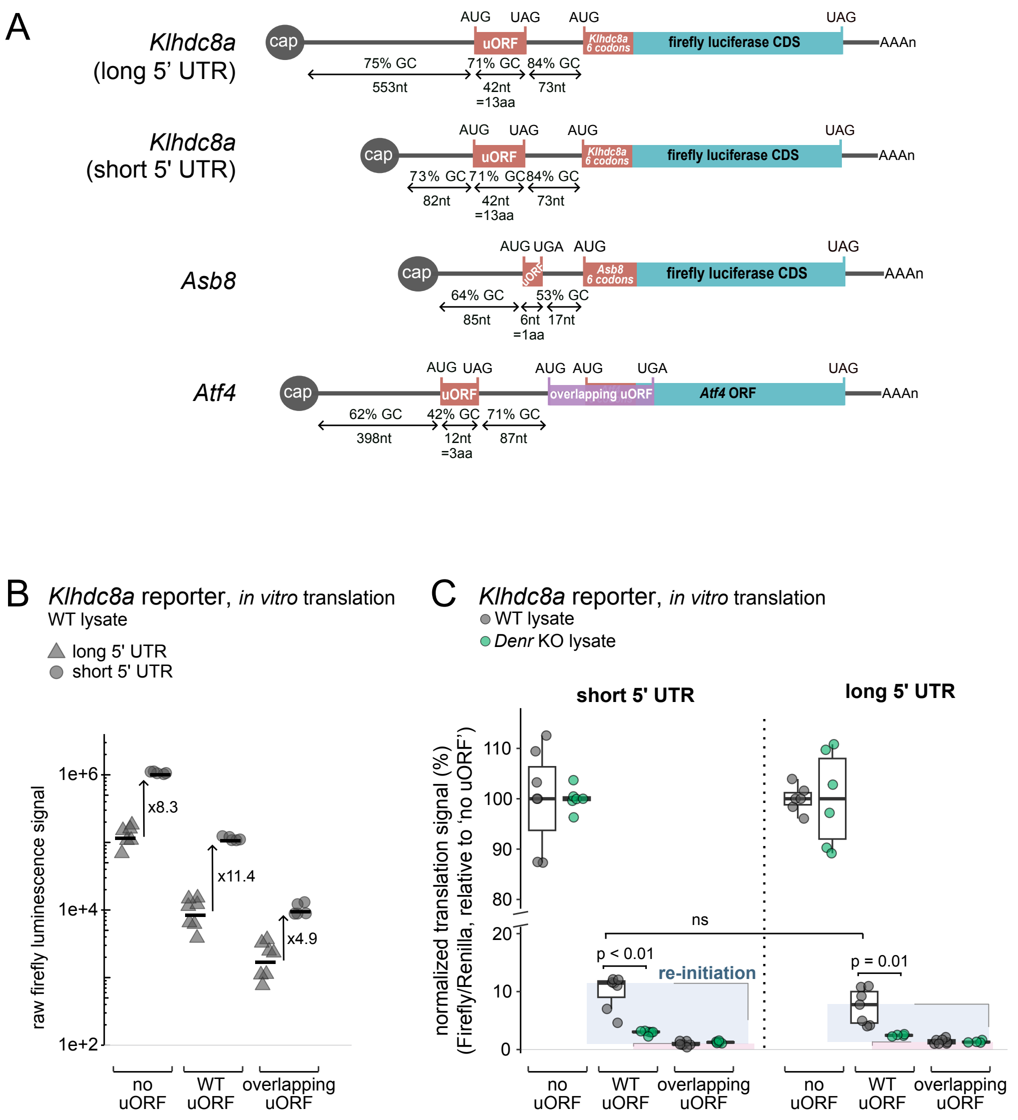
A
Klhdc8a
(long 5' UTR)
cap
AUG UAG AUG UAG
uORF
Klhdc8a 6 codons
firefly luciferase CDS
AAAn
75% GC 71% GC 84% GC
553nt 42nt =13aa 73nt
Klhdc8a
(short 5' UTR)
73% GC 71% GC 84% GC
82nt 42nt =13aa 73nt
Asb8
AUG UGA AUG UAG
uORF
Asb8 6 codons
64% GC 53% GC
85nt 6nt =1aa 17nt
Atf4
AUG UAG AUG AUG UGA UAG
overlapping uORF
Atf4 ORF
62% GC 42% GC 71% GC
398nt 12nt =3aa 87nt
B
Klhdc8a reporter, in vitro translation
WT lysate
long 5' UTR
short 5' UTR
raw firefly luminescence signal
1e+6
1e+4
1e+2
x8.3
x11.4
x4.9
no uORF
WT uORF
overlapping uORF
C
Klhdc8a reporter, in vitro translation
WT lysate
Denr KO lysate
short 5' UTR
long 5' UTR
normalized translation signal (%)
(Firefly/Renilla, relative to 'no uORF')
110
100
90
80
20
10
0
ns
p < 0.01
p = 0.01
re-initiation
no uORF
WT uORF
overlapping uORF
no uORF
WT uORF
overlapping uORF

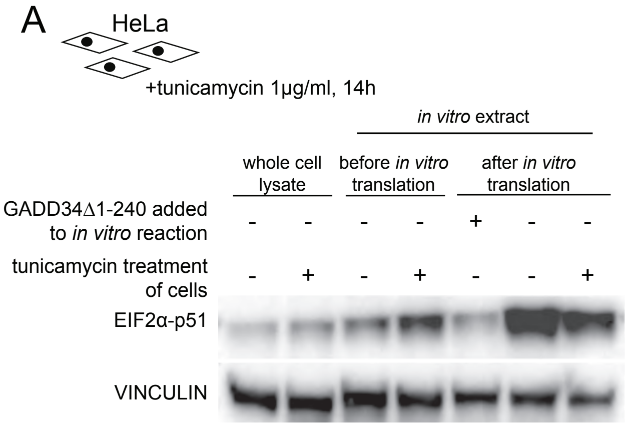
A
HeLa
+tunicamycin 1µg/ml, 14h
in vitro extract
whole cell lysate
before in vitro translation
after in vitro translation
GADD34Δ1-240 added to in vitro reaction
tunicamycin treatment of cells
EIF2α-p51
VINCULIN

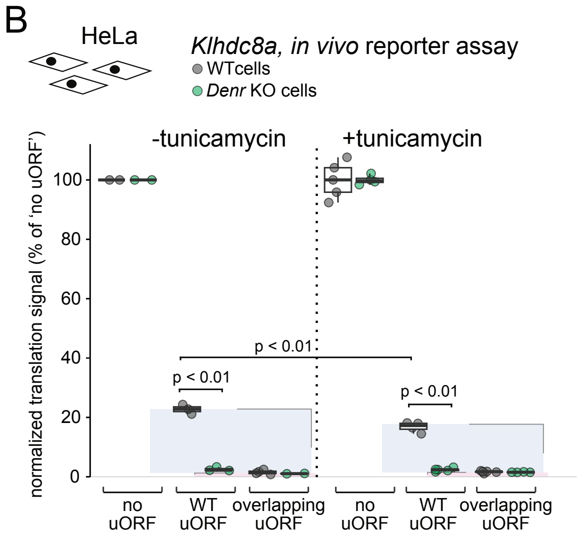
B
HeLa
Klhdc8a, in vivo reporter assay
WTcells
Denr KO cells
-tunicamycin
+tunicamycin
normalized translation signal (% of 'no uORF')
p < 0.01
p < 0.01
p < 0.01
no uORF
WT uORF
overlapping uORF

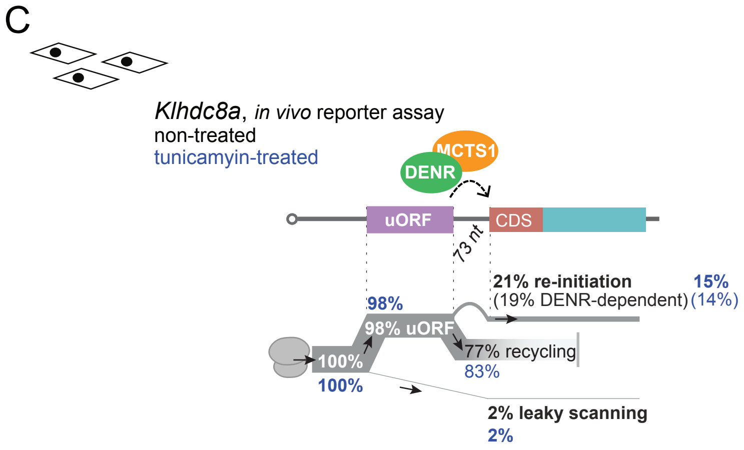
C
Klhdc8a, in vivo reporter assay
non-treated
tunicamyin-treated
MCTS1
DENR
uORF
CDS
73 nt
21% re-initiation
(19% DENR-dependent)
15%
(14%)
98%
98% uORF
77% recycling
83%
100%
100%
2% leaky scanning
2%

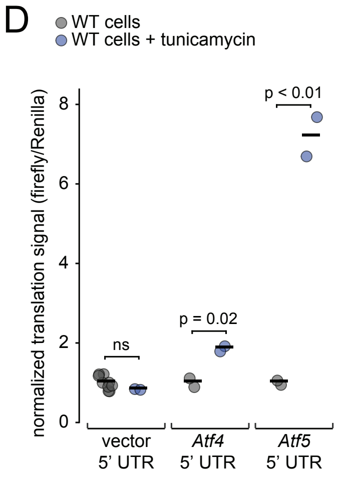
D
WT cells
WT cells + tunicamycin
normalized translation signal (firefly/Renilla)
p < 0.01
p = 0.02
ns
vector 5' UTR
Atf4 5' UTR
Atf5 5' UTR

Meurs et al., Supplementary Figure S4

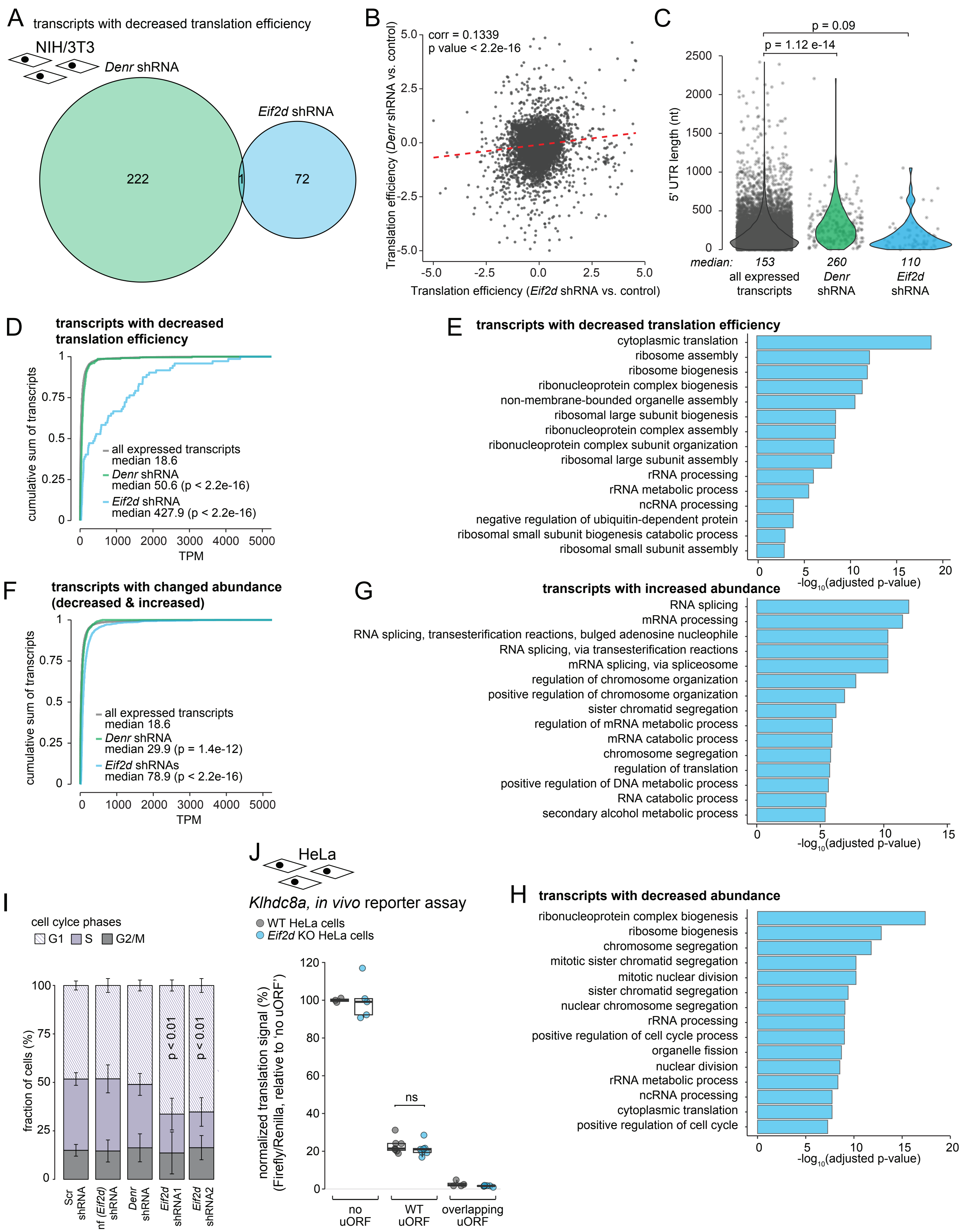

A

Alignment of mouse *Mcts1* and *Mcts2* coding sequences:

```
MmMcts1_CDS      ATGTTCAAGAAATTTGATGAAAAAGAAAATGTGTCCAACTGCATCCAGTTGAAAACCTCG      60
MmMcts2_CDS      ATGTTCAAGAAATTTGACGAGAAGGAAAGTGTGTCCAACTGCATCCAACTGAAAACTTCC      60
                 ***************** ** ** **** ******************  ******* **

MmMcts1_CDS      GTTATTAAGGGTATTAAAAATCAATTGCTAGAGCAATTTCCAGGTATTGAACCATGGCTT      120
MmMcts2_CDS      GTTATTAAGGGTATTAAGAGCCAACTGACTGAGCAGTTTCCAGGTATCGAGCCGTGGCTT      120
                 ***************** *  *** **   ***** *********** ** ** ******

MmMcts1_CDS      AATCAAATCATGCCTAAGAAAGACCCTGTGAAAATTGTCCGATGCCATGAACACATAGAA      180
MmMcts2_CDS      AATCAAATCATGCCTAAGAAAGATCCCGTCAAAATAGTGAGATGCCATGAACACATGGAA      180
                 *********************** ** ** ***** **  **************** ***

MmMcts1_CDS      ATCCTTACAGTAAATGGAGAATTACTGTTTTTTAGACAAAGAGAAGGGCCTTTTTATCCA      240
MmMcts2_CDS      ATCCTTACAGTCAACGGAGAATTACTGTTTTTCAGGCAGAGAAAAGGACCTTTTTATCCA      240
                 *********** ** ***************** ** ** *** **** ************

MmMcts1_CDS      ACTTTAAGATTACTTCATAAATATCCTTTTATCTTGCCACATCAGCAGGTTGATAAAGGA      300
MmMcts2_CDS      ACGCTAAGACTACTTCACAAATACCCGTTTATCCTGCCACACCAGCAGGTCGACAAAGGA      300
                 **  ***** ******* ***** ** ****** ******* ******** ** ******

MmMcts1_CDS      GCCATCAAATTTGTACTCAGTGGAGCAAATATCATGTGTCCTGGCTTAACTTCTCCCGGA      360
MmMcts2_CDS      GCCATCAAATTTGTGCTCAGTGGTGCAAATATCATGTGCCCGGGTTTAACGTCTCCTGGA      360
                 ************** ******** ************** ** ** ***** ***** ***

MmMcts1_CDS      GCTAAGCTTTATCCTGCTGCAGTAGATACTATTGTTGCAATCATGGCAGAAGGAAAACAA      420
MmMcts2_CDS      GCGAAGCTCTACACTGCTGCAGTAGATACCATCGTGGCGGTCATGGCAGAGGGGAAAGAG      420
                 ** ***** **  **************** ** ** **  ********** ** *** *

MmMcts1_CDS      CATGCTTTATGTGTGGGTGTCATGAAGATGTCTGCAGAAGATATTGAGAAAGTAAACAAA      480
MmMcts2_CDS      CATGCCCTGTGTGTCGGAGTCATGAAGATGGCTGCAGCAGACATTGAGAAAATCAACAAG      480
                 *****  * ***** ** ************ ****** *** ********* * *****

MmMcts1_CDS      GGAATTGGCATTGAAAATATCCATTATCTAAATGATGGTCTGTGGCATATGAAGACATAT      540
MmMcts2_CDS      GGGATCGGCATTGAGAATATCCATTATCTAAATGACGGGCTGTGGCACATGAAGACATAT      540
                 ** ** ******** ******************** ** ******** ************

MmMcts1_CDS      AAATGA      546
MmMcts2_CDS      AAGTGA      546
                 ** ***
```

A

*Atf4* - ENSMUST00000109605

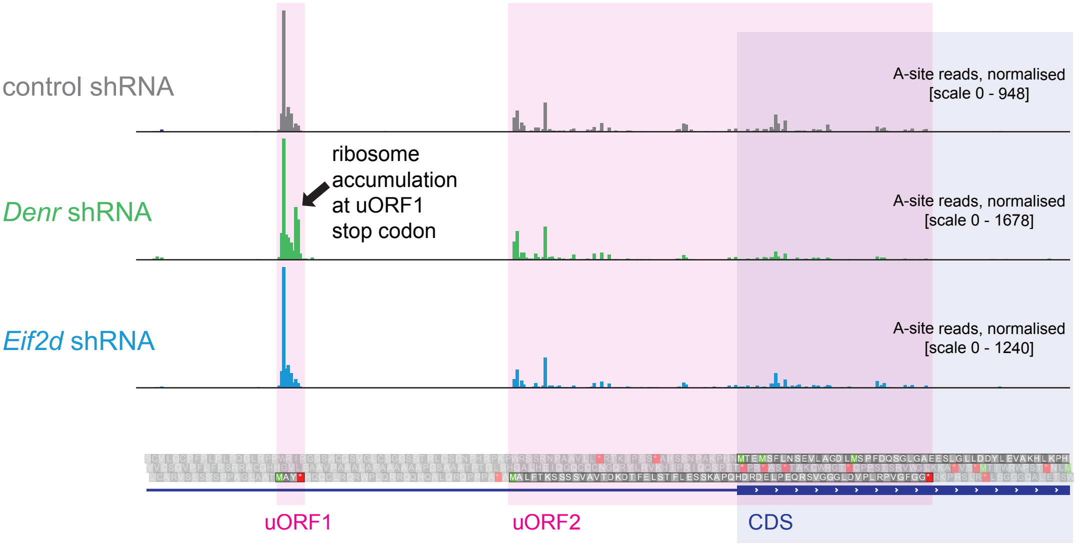

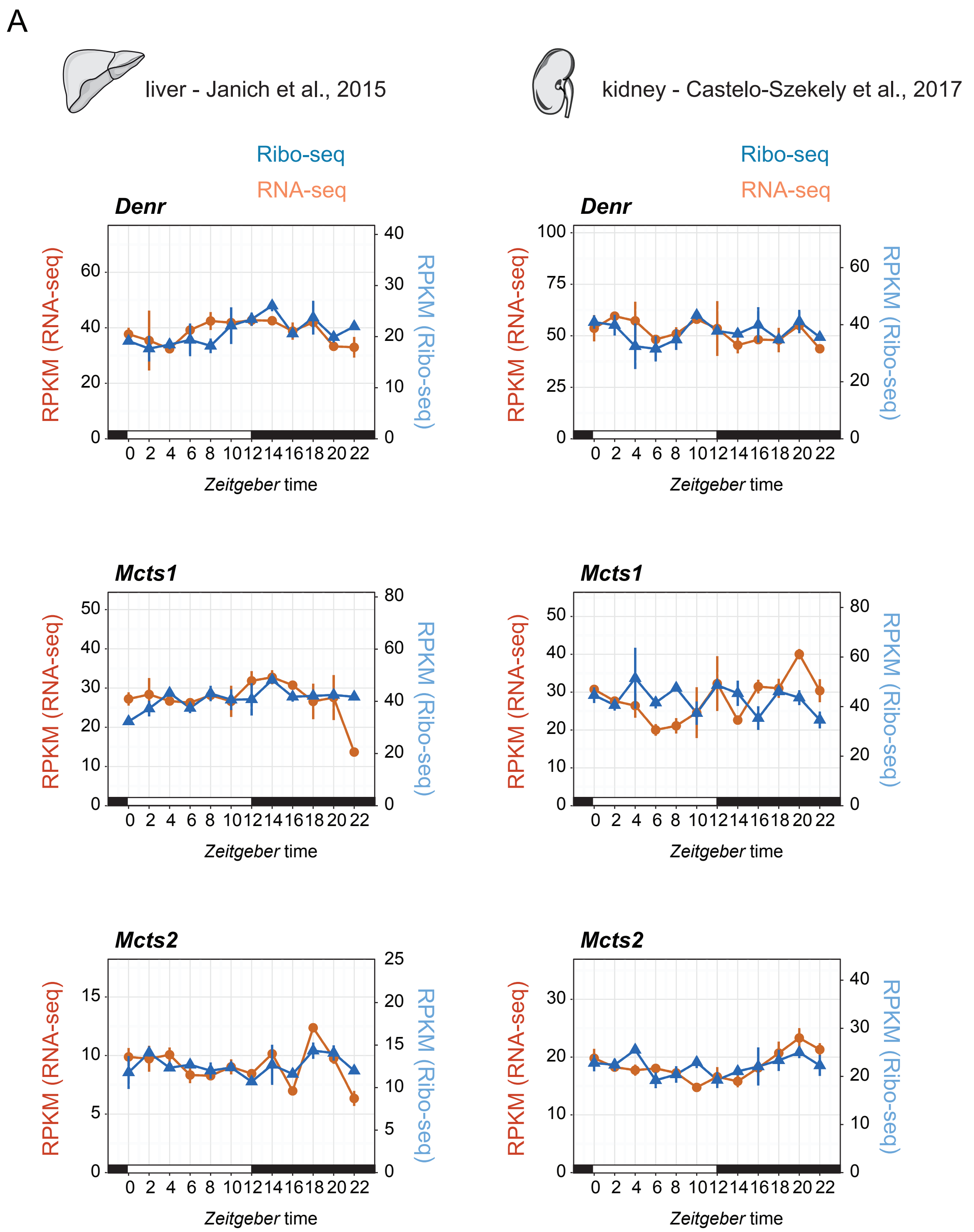
A
liver - Janich et al., 2015
kidney - Castelo-Szekely et al., 2017
Ribo-seq
RNA-seq
Denr
Mcts1
Mcts2
RPKM (RNA-seq)
RPKM (Ribo-seq)
Zeitgeber time
0 2 4 6 8 10 12 14 16 18 20 22
